# Coincidence detection of mitogenic signals via cytosolic pH regulates Cyclin D1 expression

**DOI:** 10.1101/2020.07.15.202333

**Authors:** Lisa Maria Koch, Eivind Birkeland, Stefania Battaglioni, Xiao Helle, Mayura Meerang, Stefanie Hiltbrunner, Alfredo J. Ibáñez, Matthias Peter, Alessandra Curioni, Isabelle Opitz, Reinhard Dechant

## Abstract

Enhanced cell growth and proliferation are accompanied by profound changes in cellular metabolism. Originally identified as the Warburg effect in cancer, such metabolic changes are also common under physiological conditions and include increased fermentation and elevated cytosolic pH (pHc)^1,2^. However, how these changes contribute to enhanced cell growth and proliferation is unclear. Here, we demonstrate that elevated pHc specifically orchestrates an E2F-dependent transcriptional program to drive cell proliferation by promoting Cyclin D1 expression. pHc-dependent transcription of Cyclin D1 requires the transcription factors CREB1/ATF1 and ETS1 and the Histone Acetyltransferases p300/CBP. Interestingly, biochemical characterization revealed that the CREB1-p300/CBP interaction acts as a pH-sensor and coincidence detector linking different mitotic signals to Cyclin D1 transcription. We also show that elevated pHc contributes to increased Cyclin D1 expression in Malignant Pleural Mesotheliomas (MPMs) and renders them hypersensitive to pharmacological reduction of pHc. Taken together, these data demonstrate that elevated pHc is a critical cellular signal regulating G1 progression and provide a mechanism linking elevated pHc to oncogenic activation of Cyclin D1 in MPMs and possibly other Cyclin D1-dependent tumors. Thus, an increase of pHc may represent a functionally important, early event in the etiology of cancer amenable to therapeutic intervention.

## Main

To test for the significance of pHc in the regulation of cell growth and proliferation, we established conditions to control pHc in cultured mammalian cells using inhibition of Sodium Hydrogen Exchangers (NHEs), the main regulators of pHc in most cell types. In RPE-1 cells, glucose and growth factors in the form of fetal calf serum (FCS) cooperatively increased pHc upon addition to starved cells (Figures 1a and S1a-b), suggesting that pHc is regulated both by metabolic cues and by mitogen activated kinases. Dimethyl amiloride (DMA), an NHE inhibitor^3^ and derivative of a widely used potassium-sparing diuretic, completely alleviated the observed increase in pHc (Figures 1a and S1c) and lead to decreased cell growth (Figure 1b and ^4^). Similarly, RNAi against NHE1, the most abundant NHE protein at the plasma membrane (Figure S1d), also decreased pHc and cell growth (Figures 1c-d), demonstrating that NHE inhibition can be used to modulate pHc-dependent cell growth and proliferation.

**Figure 1:**
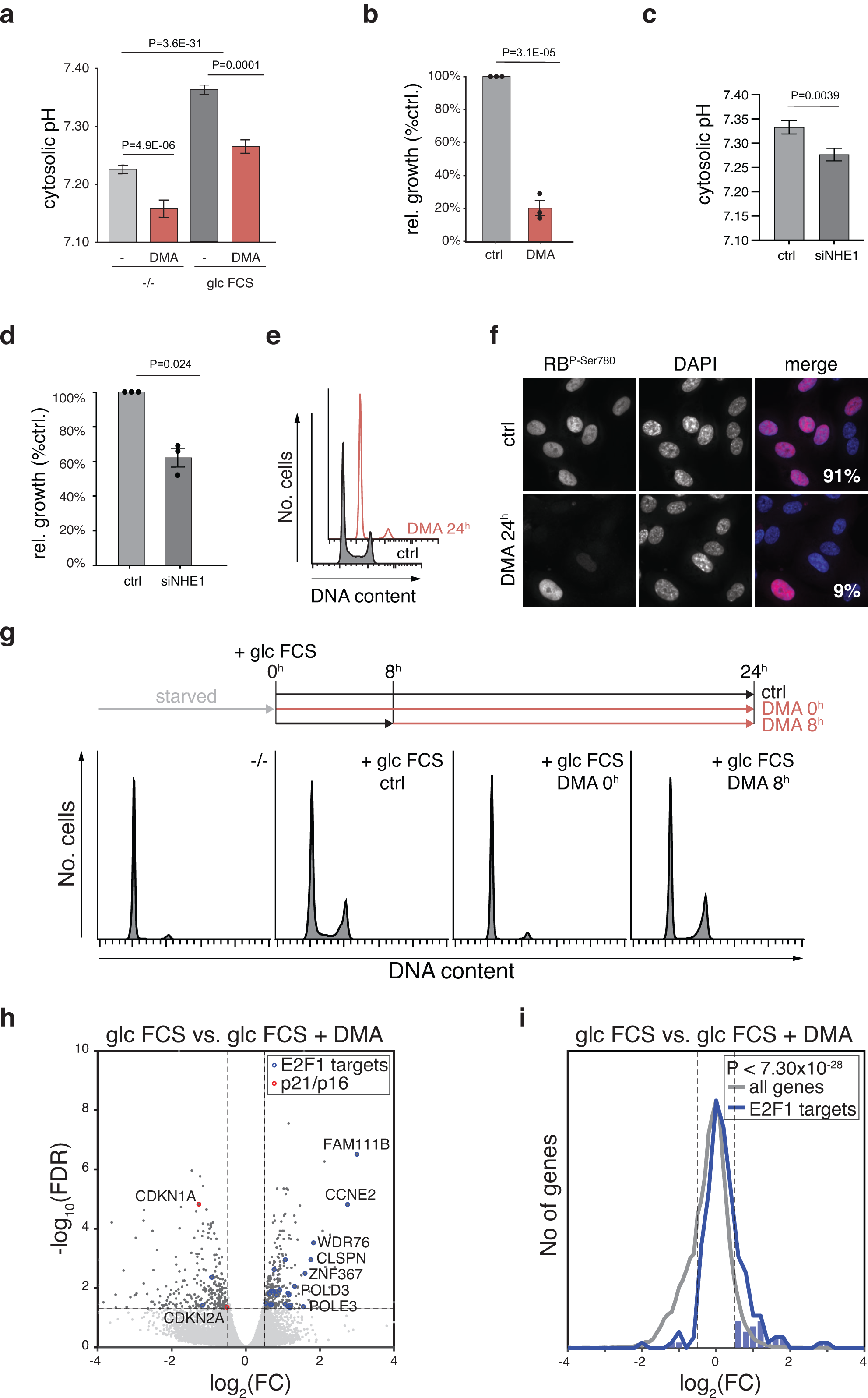
NHE1 activity regulates pHc and promotes cell cycle progression through early G1. (**a-d**) pHc and cell growth are regulated by glucose and growth factors in an NHE1 dependent manner. (**a**) RPE-1 cells expressing pHluorin were starved and pHc was determined 30 min after stimulation with the indicated conditions. (mean +/- S.E.M. of pooled single cell data of independent experiments N≧3). (**b**) Cells were grown in the presence or absence of DMA for 3 days and biomass was determined using MTT. (mean +/- S.E.M. N=3, one sample t-Test) **(c-d)** RPE-1 cells expressing pHluorin were subjected to siRNA against NHE1 and pHc **(c)** and relative biomass accumulation **(d)** was determined after 72h. (**e-f**) Elevated pHc promotes progression through early G1. DNA content (**e**) and RB phosphorylation at Ser780 (**f**) were determined in RPE-1 cells 24^h^ after treatment with DMA. Percentage (%) of cells positive for phospho-RB is indicated (mean, N=3). (**g**) RPE-1 cells were arrested in G1 by FCS and glucose starvation and DNA content was followed by flow cytometry. Cells were treated with DMA simultaneously with FCS and glucose (0^h^) or 8^h^ post stimulation as indicated. (**h** and **i**) pHc regulates G1/S progression through a transcriptional program. (**h**) Volcano-plot of genes regulated by DMA treatment. Verified E2F targets and CDK-inhibitors p16 and p21 are shown in blue and red, respectively. (**i**) Histogram-based distributions of fold-changes for all genes and verified E2F targets are shown.

### pHc promotes cell cycle progression

Interestingly, treatment of cycling cells with DMA or RNAi against NHE1 caused a pronounced arrest in G_1_ with low pRB phosphorylation (Figures 1e-f; S1e-g), suggesting that an elevated pHc is particularly important for CDK activation. Identical phenotypes as in hTERT immortalized RPE-1 cells were observed in primary fibroblasts (HFF-1) and different cancer cell lines (Figure S2), or upon reduction of pHc by reducing extracellular pH (Figure S1h). Importantly, pHc does not indirectly regulate proliferation by perturbing energy or central carbon metabolism (Figure S3). Rather, pHc appears to generally regulate a specific step in G_1_ to signal cell cycle progression.^5,6^

Similarly, while cells arrested in G_1_/G_0_ by starvation readily progressed through S phase upon re-addition of FCS and glucose, DMA completely prevented cell cycle progression and largely abolished RB phosphorylation and Cyclin A expression when added simultaneously with the mitogenic stimulus (Figures 1g and S4a). In contrast, addition of DMA 8^h^ after glucose and FCS addition, but before detection of S-phase initiation by BrdU incorporation (Figure S4b) did not affect the number of cells in G2 or M phase 24^h^ post stimulation or Cyclin A expression (Figures 1g and S4a), consistent with previous results^5^. As expected, stimulation of starved cells with FCS and glucose triggered a global change in gene expression including genes required for ribosome biogenesis and cell cycle control (Figure S4c and ^7,8^). DMA treatment revealed a similar transcriptional response compared with starvation (Figures S4d-e). However, DMA treatment mostly impaired expression of genes required for cell cycle progression, but had little effect on ribosome biogenesis (Figure S4c). Thus, although DMA treatment mimics starvation with respect to cell cycle and growth arrest, an elevated pHc triggers a specific transcriptional program to promote proliferation and may only indirectly affect cell growth. In particular, a large number of genes regulated by E2F, the main transcriptional regulator of G1/S progression, were dependent on elevated pHc for expression, while expression of Cyclin-dependent kinase inhibitors p16 and p21 was induced upon DMA treatment (Figures 1h-i). Thus, an elevated pHc is necessary to specifically promote progression through early G_1_, possibly by regulating CDK4^Cyclin D1^ expression and/or activity (Figure S4b, ^7,9^)

Indeed, treatment of cells with DMA delayed accumulation of Cyclin D1 protein and prevented the increase in Cyclin D1 mRNA upon mitogenic stimulation in different cell lines (Figures 2a-b and S5a). DMA treatment or RNAi against NHE1 also reduced the abundance of Cyclin D1 mRNA and protein in cycling cells (Figure 2c and S5b-c). Inhibition of Cdk4^Cyclin D1^ activity using Ribociclib^10^ prevented cell cycle reentry and RB phosphorylation also when added 8^h^ post mitogenic stimulation (Figures S5d and e). Thus, high pHc drives cell cycle progression through early G1 by regulating Cyclin D1 mRNA levels. Once Cyclin D1 levels have reached a certain threshold necessary to pass the restriction point,^7,9^ pH-dependent signaling is dispensable for CDK4^Cyclin D1^ activity and further cell cycle progression.

**Figure 2:**
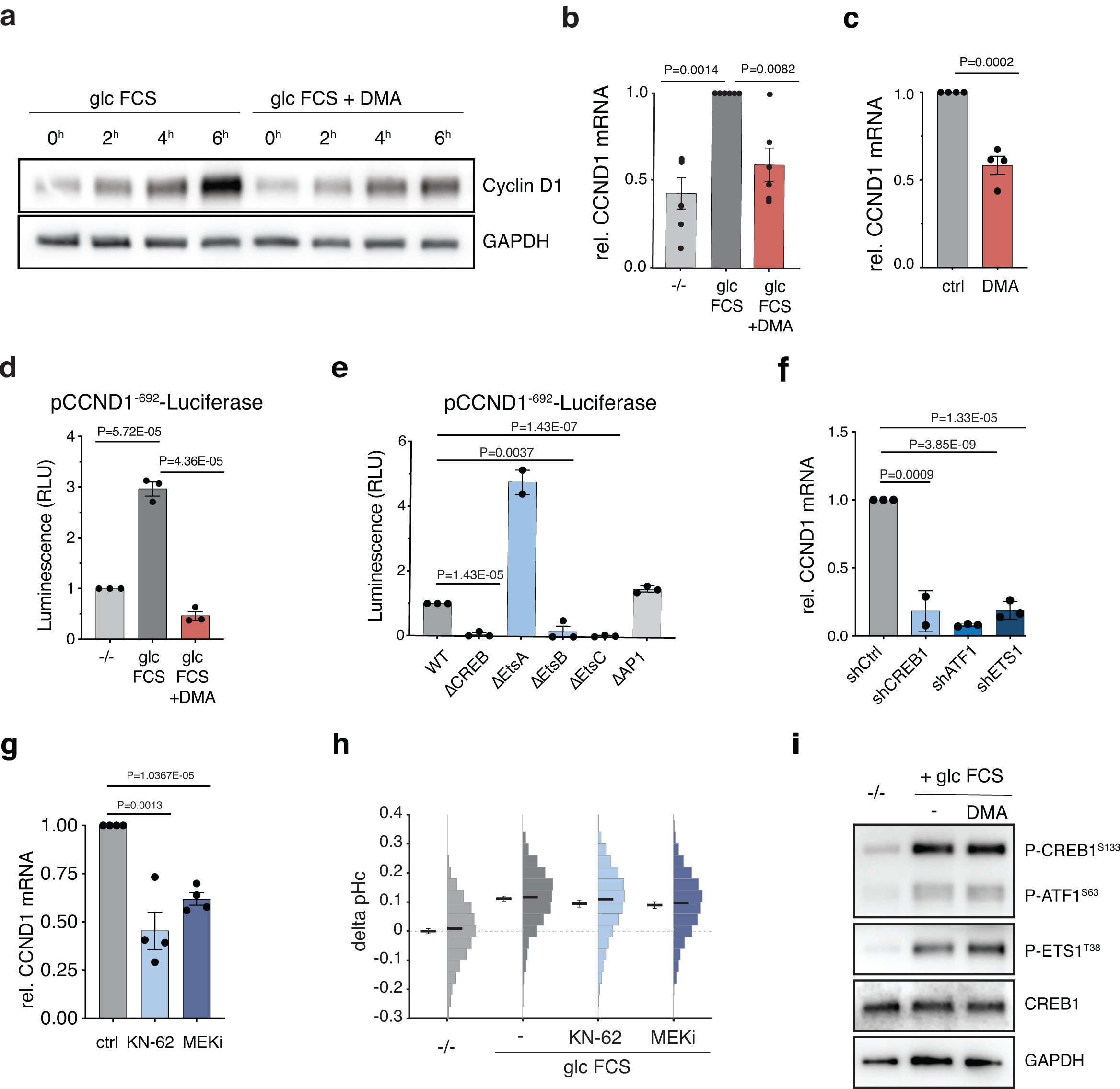
High pHc regulates Cyclin D1 transcription in parallel or downstream of CREB activation. (**a**-**c**) pHc regulates Cyclin D1 protein and mRNA abundance. RPE-1 cells were arrested in G1 by starvation and levels of Cyclin D1 protein (**a**) and mRNA (**b**) were determined upon stimulation with FCS and glucose at the indicated time points (4h for qPCR) with or without DMA by western blotting and qPCR, respectively. (mean +/- S.E.M. N=4, one sample t-Test) (**c**) RPE-1 cells were grown in RPMI media and treated with DMA for 4h and Cyclin D1 mRNA levels were determined. (mean +/- S.E.M. N=3, one sample t-Test) (**d-f**) pHc regulates Cyclin D1 transcription in a CREB- family and ETS1-dependent manner. (**d**) RPE-1 cells expressing pCCND1^692^-Luciferase were starved and Luciferase activity was measured 6 h after stimulation with the indicated conditions. A result representative of three independent experiments is shown (mean +/- S.E.M. N=3 technical replicates) (**e**) Luciferase activity of RPE-1 cells expressing wild-type or mutated reporter pCCND1^692^-Luciferase was determined. (mean +/- S.E.M. N=3) (**f**) Cyclin D1 mRNA abundance was determined in RPE-1 cells upon shRNA mediated knock-down of the indicated genes. (mean +/- S.E.M. N=3) (**g- h**) pHc regulates Cyclin D1 expression downstream or in parallel to CREB-family and ETS1 activation. (**g**) RPE-1 cells were treated with MEK or CamKIV inhibitors and Cyclin D1 (**g**) mRNA levels (mean +/- S.E.M. N=4) and (**h**) pHc were determined. (mean +/- S.E.M. of pooled single cell data, N=3) (**i**) Starved RPE-1 cells were stimulated with glucose and FCS in the presence or absence of DMA and phosphorylation of CREB1, ATF1 and ETS1 was followed by western blotting.

### pHc regulates Cyclin D1 transcription

A luciferase reporter construct under the control of the Cyclin D1 promoter^11^ fully replicated pHc-dependent regulation of the endogenous Cyclin D1 mRNA (Figure 2d), suggesting that pHc regulates Cyclin D1 transcription rather than mRNA stability. Likewise, inhibition of Pol-II dependent transcription using Actinomycin D decreased Cyclin D1 mRNA, but was not additive with DMA treatment (Figure S5f).

Therefore, we sought to identify transcription factors mediating pHc-dependent Cyclin D1 transcription. Interestingly, deleting the CREB and the EtsB binding sites led to strongly reduced expression of the Cyclin D1 luciferase reporter (Figure 2e and ^11^). Similarly, knock-down of CREB1 and ATF1 as well as ETS1 reduced Cyclin D1 mRNA and protein levels (Figures 2f and S5g), suggesting that at least two functionally distinct inputs are simultaneously required to regulate Cyclin D1 expression.

CREB and ETS family transcription factors are regulated by various upstream signals including PKA, CamKIV and MAPK signaling.^12-14^ Inhibition of MEK1 and CamKIV reduced Cyclin D1 mRNA without affecting pHc (Figures 2g-h). However, DMA treatment did not reduce Ets1-T38, Creb1-S133 or Atf1-S63 phosphorylation, sites critically required for transcriptional activation of their downstream targets^15^ (Figure 2i). Therefore, pHc-dependent activation of Cyclin D1 transcription acts downstream of, or in parallel to, CREB1/ATF1 and ETS1 activation.

CREB and ETS family transcription factors recruit the transcriptional co-activators p300/CBP, two histone acetyltransferases (HATs) that catalyze the acetylation of Histone H3-K27 and transcriptional activation of their target genes^16-18^. Knock-down of CREB1, ATF1 and ETS1 led to a global reduction of H3-K27 acetylation (Figure S6a), suggesting that p300/CBP may mediate pHc-dependent transcription. Indeed, reduced activity of p300/CBP mediated by shRNA or by the pharmacological inhibitor C646 decreased Cyclin D1 expression and H3-K27 acetylation (Figure 3a-b and S6a-c). Treatment of cells with DMA also globally reduced H3-K27 (Figure 3b), but not H3-K9 acetylation (Figure S5d), suggesting that reduced pHc does not generally prevent histone acetylation, but regulates a specific p300/CBP-dependent transcriptional program, similar to previously published data.^19^ Indeed, we found a striking overlap between genes regulated by starvation, DMA and C646 treatment (Figure 3c and S7). Co-regulated genes were highly enriched for targets of E2F, ETS1 and CREB1, further confirming the importance of these transcription factors in orchestrating pHc-dependent cell cycle progression (Supplementary Table S15).

**Figure 3:**
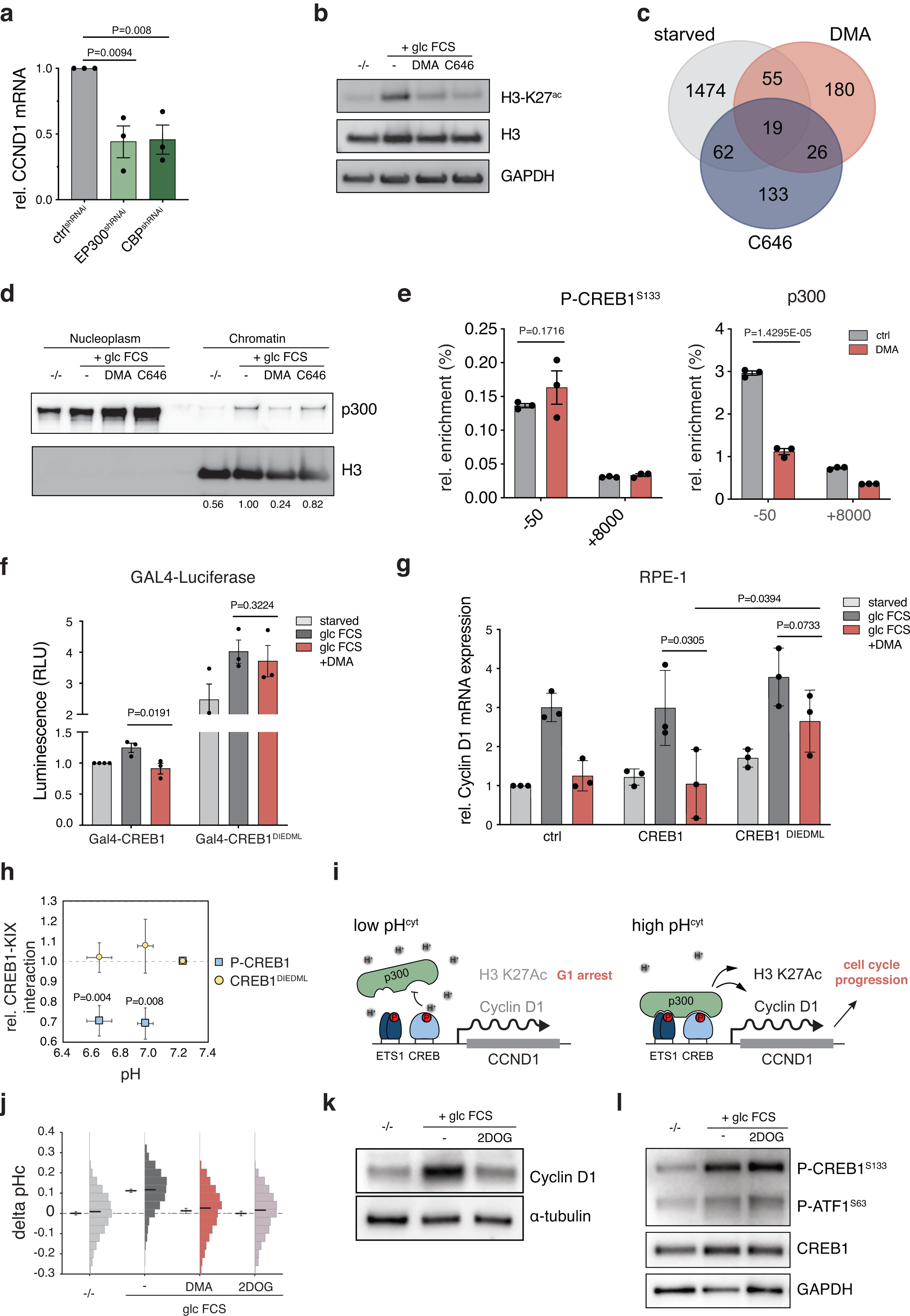
Sensing of pHc by the CREB/p300 interaction regulates Cyclin D1 expression. (**a-c**) pH-dependent p300/CBP recruitment to chromatin regulates transcription. (**a**) Cyclin D1 mRNA levels upon p300 and CBP knockdown in RPE-1 cells was determined by qPCR. (mean +/- S.E.M. N=3, one sample t-Test) (**b**) RPE-1 cells were starved, stimulated with FCS and glucose in the presence of DMA or the p300/CBP inhibitor C646 as indicated and H3-K27 acetylation was analyzed by western blotting. (**c**) RPE-1 cells were arrested in G1 by serum and glucose starvation, stimulated with FCS and glucose with or without DMA or C646 for 4h and subjected to RNAseq. Venn diagram highlighting the similarity of changes in the transcriptome under all treatments. Only significantly repressed genes relative to the control condition are shown. (**d**) Extracts from RPE-1 cells treated with DMA and C646 were subjected to subcellular protein fractionation and chromatin recruitment of p300 was determined by western blotting. Mean of relative p300 recruitment is indicated (N=5) (**e**) Recruitment of p300, but not phosphorylated CREB1, to the Cyclin D1 promoter is regulated by pHc. RPE-1 cells treated with DMA for 4h were subjected to ChIP analysis. Binding of the indicated proteins to the CREB-family binding site within the promoter (−50) and within intron 4 (+8000 nt) is shown. (Result is representative of three independent experiments) (**f-g**) CREB1 phosphorylation mediates pH-dependent transcriptional activation. (**f**) RPE-1 cells expressing pGAL4-Luciferase and Gal4-CREB1 constructs as indicated were starved and luciferase activity was determined upon stimulation of FCS and glucose with or without DMA. (mean +/- S.E.M. N=3) (**g**) RPE-1 cells stably expressing CREB1-WT or mutant constructs as indicated were starved and Cyclin D1 mRNA abundance was determined upon stimulation of FCS and glucose with or without DMA. (mean +/- S.E.M. N=3) (**h**) CREB1 phosphorylation mediates pH-dependent binding to p300/CBP *in vitro*. Biotinylated peptides corresponding to phosphorylated wild-type CREB1 or mutant CREB1 were bound to Streptavidin beads and pH-dependence of binding of the CBP KIX domain was determined. (mean +/- S.E.M. N=6) (**i**) Schematic representation of the proposed model for regulation of Cyclin D1. (**j-l**) Glucose metabolism mediates Cyclin D1 expression via pHc. RPE-1 cells were starved and stimulated with FCS and glucose in the presence of 2DOG and DMA and (**j**) changes in cytosolic pH (mean +/- S.E.M. of pooled single cell data, N=3), (**k**) Cyclin D1 expression and (**l**) CREB1 phosphorylation were determined by western blotting.

Taken together, these data suggest that pHc may regulate p300/CBP activity and/or chromatin recruitment. Subcellular protein fractionation assays revealed strong recruitment of p300 to chromatin upon mitogenic stimulation, which was lost upon DMA treatment (Figure 3d). Although detection of phosphorylated CREB1-S133 at the Cyclin D1 promoter by ChIP was not diminished by DMA treatment, cells treated with DMA significantly reduced recruitment of p300 to the Cyclin D1 promoter (Figure 3e). Thus, pHc may regulate recruitment of p300/CBP to the Cyclin D1 promoter by directly modulating binding of p300/CBP to CREB1/ATF1 and/or ETS1.

### The CREB1/p300 interaction acts as a pH sensor

To test this hypothesis, we employed a luciferase reporter based on a synthetic Gal4-dependent promoter and expressed CREB1 as a Gal4 fusion protein. Expression of wild-type Gal4-CREB1 retained DMA-sensitive reporter expression (Figure 3f), further supporting that pHc does not regulate CREB1 activation or DNA binding. p300/CBP interacts with the KID domain of CREB1, which shares some sequence similarity with the transcription factor SREBP that does not require phosphorylation for p300/CBP binding.^15^ Replacing 6 amino acids of CREB1 encompassing Ser133 with a stretch of mostly acidic residues from SREBP leads to constitutive and strong CREB-p300 interaction^15,20^ and the resulting construct (Gal4-CREB1^DIEDML^) rendered luciferase expression insensitive to DMA (Figure 3f). Expression of full-length CREB1^DIEDML^ also partially rescued Cyclin D1 expression, cell growth and RB phosphorylation in the presence of DMA (Figure 3g and S8a-b). Thus, pHc does not affect p300/CBP activity, but modulates the CREB1-p300/CBP interaction.

Thus, we sought to reconstitute the interaction between CREB1 and CBP *in vitro*. As expected, a short phosphorylated CREB1 peptide (biotin-P-CREB1^121-151^) is sufficient to confer binding to the KIX domain of CBP (residues 586-672)^21^ (Figure S8b). Importantly, the interaction of phosphorylated wild-type CREB1^121-151^, but not of CREB1^121-151^-DIEDML with the KIX domain was significantly impaired when reducing pH within the physiological range (Figure 3h).

Within the residues of CREB1 necessary for binding to the KIX domain only the side chain of phosphoserine, with a known pKa of 5.78 for isolated phosphoserine,^22^ can change its protonation state close to the physiological pH range, suggesting that pH-sensitivity is mediated by the phospho-serine itself. In contrast, the hyperactive DIEDML mutant introduces 3 acidic residues likely acting as a phosphomimetic, pH-insensitive mutant. We conclude that the P-CREB1-p300/CBP interaction acts as a pH sensor and establishes a coincidence detector for two distinct mitogenic signals, CREB1 activation by phosphorylation and elevated pHc, both of which are necessary to allow efficient Cyclin D1 expression (Figure 3i).

Mechanisms explaining the FCS-dependent increase of pHc may involve MAPK or Akt-dependent phosphorylation of NHE1 at Ser770, Ser771 and Ser648, respectively.^23^ Yet, although inhibition of Akt, but not MAPK reduced pHc (Figures 2h and S8a), and Akt activity was independent of pHc (Figure S8d), we failed to detect phosphorylation of NHE1 at Ser648 *in vivo* (data not shown). Instead, Akt may promote pHc via stimulation of glycolytic activity.^24^ Consistently, inhibition of glycolysis by 2-deoxyglucose prevents Cyclin D1 expression and reduces pHc without affecting CREB1 phosphorylation (Figure 3k-l), suggesting that pHc may represent a glucose-dependent, metabolic input into cell proliferation.

### pHc is increased in MPMs

pHc-dependent Cyclin D1 expression could also readily explain the oncogenic potential of elevated pHc.^25^ Mechanisms underlying elevated pHc may vary in different tumors, but might include overexpression of NHE1 or stimulation of specific NHE1 activity. In highly glycolytic tumors, such as pancreatic ductal adenocarcinoma (PDAC), high rates of aerobic fermentation may promote the specific activity of NHE1 and, thus, upregulate Cyclin D1 in a pHc-dependent manner. Indeed, in PDAC cell lines, inhibition of NHE- and Akt-activity reduced pHc (Figures S9a). Inhibition of NHE activity by DMA or siRNA mediated knock-down also lead to cell cycle arrest with low RB phosphorylation and Cyclin D1 expression (Figure S2 and S9b-c), consistent with previous results.^26,27^ Thus, an elevated pHc in PDACs may contribute to enhanced proliferation. Yet, although Cyclin D1 expression strongly correlated with survival in PDAC patients, no such correlation was observed with NHE1 expression (Figure S9d).

In contrast, screening the CCLE database^28^ revealed a high correlation between NHE1 and Cyclin D1 mRNA levels for cell-lines derived from MPMs, an aggressive malignancy derived from the pleural epithelium functionally linked to Cyclin D1 activation (Figure 4a and S10a ^29-31^). Similarly, in MPM patients the pH of the pleural fluid has predictive potential for survival, with lower pH due to high proton extrusion from tumor cells being associated with poor prognosis^32^. Thus, NHE1 overexpression may cause Cyclin D1 up-regulation in these tumors through elevated pHc.

**Figure 4:**
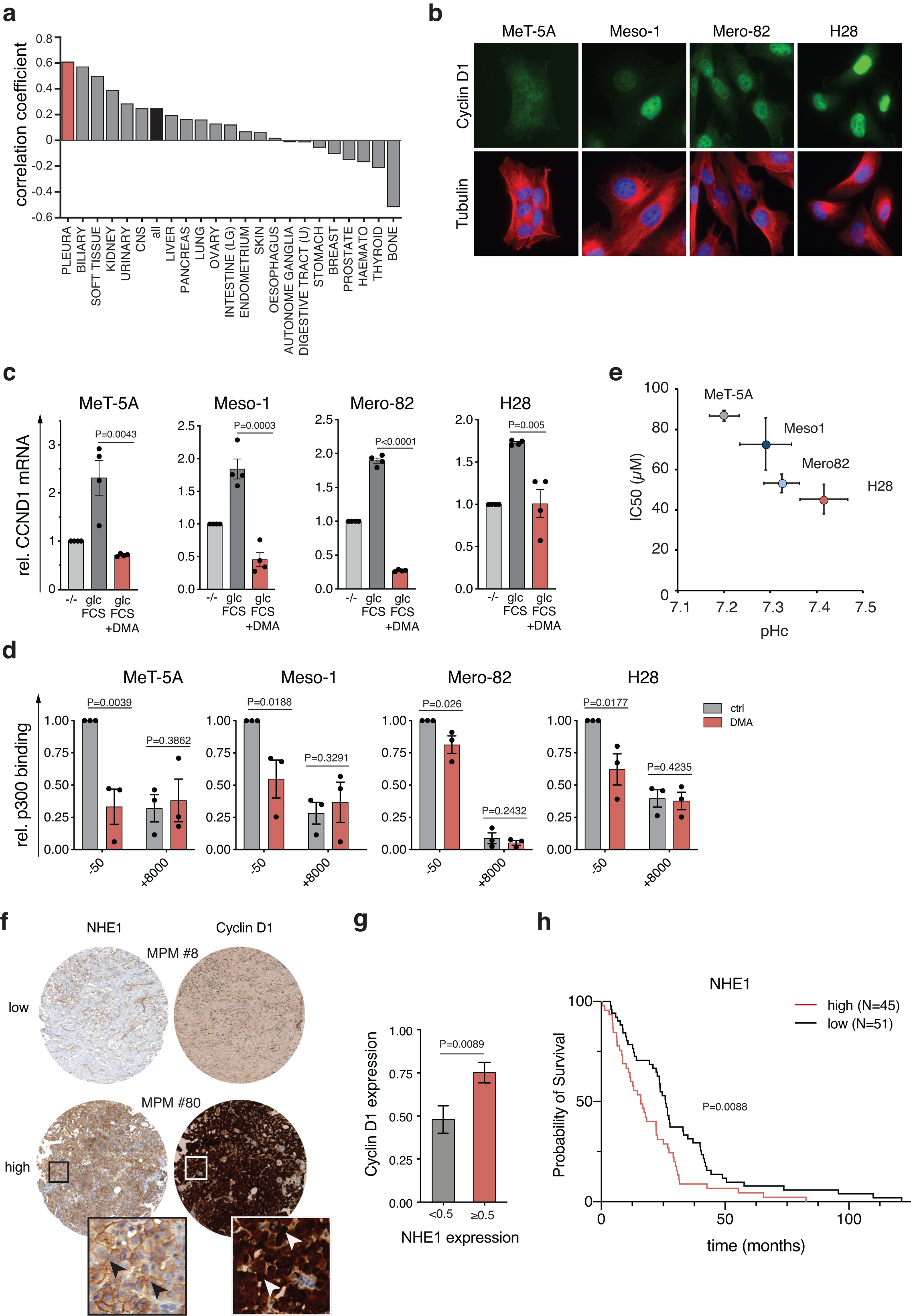
Elevated pHc mediates oncogenic activation of Cyclin D1 in MPMs. (**a**) Correlation of NHE1 and Cyclin D1 expression in different cancers. Expression levels (as RPKM) of NHE1 and Cyclin D1 were obtained from CCLE and analyzed for the individual cancer types or pooled data (all) from all cancers by using a linear fit model. Pearson correlation coefficients are shown. (**b**) Cyclin D1 expression is increased in MPM cell lines. Representative images of control cells (MeT-5A) and MPM cell lines stained for Cyclin D1 are shown (Meso1 = ACC-Meso-1). (**c-d**) MPM cell lines retain pHc-dependent cell cycle progression (**c)** MPM cell lines were starved and Cyclin D1 mRNA abundance was analyzed upon stimulation with FCS and glucose in the presence or absence of DMA. (mean +/- S.E.M. N=3) (**d**) MPM cell lines were treated with DMA for 4 hours and relative p300 recruitment to the Cyclin D1 promoter (−50) or within intron 4 (+8000 nt) was determined (mean +/- S.E.M. N=3). (**e**) The IC50 for DMA correlates with pHc in MPM cell lines. Scatter plot of the IC50s of the indicated cell lines for DMA are shown (mean +/- S.E.M. N=5) relative to steady state pHc in the presence of FCS and glucose. (mean +/- S.E.M. of pooled single cell data, N=3) (**f-h**) NHE1 expression and Cyclin D1 expression correlate in tumor sections. (**f**) representative images of histological sections stained for the indicated markers. (**g**) Relative expression of Cyclin D1 and NHE1 was scored in the different tumor sections and sections were grouped according to NHE1 expression (high NHE1 N=45, low NHE1 N=51). Mean relative Cyclin D1 expression was plotted for high and low expressing tumors (data are shown as mean +/- SEM). (**h**) Kaplan Meier analysis of the survival of patients with high and low NHE1 expression is shown.

Cell-lines derived from MPM patients had high levels of nuclear Cyclin D1 as compared to a control cell-line derived from a healthy donor (Figure 4b). In all cell-lines, Cyclin D1 expression, p300 chromatin recruitment and RB phosphorylation was sensitive to DMA (Figure 4c-d and S10b-c) or knock down of NHE1 (Figure S10d), the predominant plasma membrane NHE isoform in these cells (Figure S10e). Thus, MPM cell-lines retained pHc-dependent regulation of Cyclin D1 transcription. Steady state levels of Cyclin D1 expression correlated completely with pHc and cell-lines with higher pHc also were significantly more sensitive to inhibition by DMA (Figure 4b and e), suggesting that high activity of the pHc/Cyclin D1 axis is critical for cell proliferation in these tumors and may contribute to in initiation or progression of MPMs.

In support of this, immunohistochemical analysis of MPM biopsies showed significantly higher Cyclin D1 expression in patients with high NHE1 expression (Figure 4f-g, S11a). Finally, high NHE1 expression negatively correlates with patient survival and Cyclin D1 expression in MPMs at the level of protein and mRNA expression (Figure 4h and S11b-c). Thus, upregulation of NHE activity may be a critical step in tumor initiation and/or progression in different tumor types leading to enhanced Cyclin D1 expression.

## Discussion

Taken together, our data demonstrate that elevated pHc signals passage of the restriction point by regulating Cyclin D1 expression and contributes to tumor development. Passage of the restriction point and, thus, commitment to enter a new cell cycle requires integrating multiple environmental signals to ensure successful completion of cell division. Our data suggest that pHc- and phosphorylation-dependent recruitment of the transcriptional coactivators p300/CBP to the Cyclin D1 promoter establishes a coincidence detector for several mitogenic stimuli necessary for Cyclin D1 transcription. While phosphorylation of transcription factors is mediated by mitogenic kinases, elevated pHc is required for binding of these transcription factors to p300/CBP by modulating the protonation state of the phosphorylated residues.

Interestingly, pHc itself is regulated by growth factors and requires active glucose metabolism. While glucose stimulation might simply provide sufficient energy necessary to establish high membrane potential and NHE activity, growth factors may increase pHc by further stimulating glycolytic flux.^33^ Thus, we propose that pHc represents a metabolic signal reporting on the energy status of the cell, which is conserved from yeast to man^34,35^ and ensures initiation of cell proliferation by growth factors only in the presence of sufficient nutrient supply. This model is also reminiscent of coincidence detection by multivalent recruitment of effector proteins regulating endocytic trafficking,^36^ which likewise depend on pH-sensitive interactions with phosphoinositides.^37^ Thus, pHc may be a common modulator of phospho-dependent interactions coordinating a variety of cellular processes.^19,38^

This model also provides a molecular explanation for how increased pHc, which has long been recognized as a hallmark of cancer,^25,39-41^ can contribute to cellular transformation and cancer progression.^42,43^ Any genetic alteration leading to an elevated pHc may help to evade growth suppression by stimulating a transcriptional program driving G1 progression and thus may represent an early step in cellular transformation that can be a target of pharmacological intervention.^44^

## Materials and Methods

A detailed description of materials and methods used can be found in the supplemental information.

## Author contributions

RD conceptualized the study; performed formal analysis and visualization; wrote the manuscript with contributions of all authors; LMK, EB, SB, XH and RD performed experiments; LMK, EB, SH and RD analyzed data; AJI performed metabolomics analysis; RD, IO, AC and MP supervised the study.

## Acknowledgements

We thank Philipp Kimmig, Samuel Gilberto, Werner Kovacs, Markus Stoffel, Alicia Smith and members of the Peter, Curioni and Opitz laboratories for helpful discussions and comments on the manuscript; Bart Vrugt, Christiane Mittmann and Marcel Glönkler, Department of Pathology and Molecular Pathology, University Hospital Zurich, for help with establishment of NHE1 and Cyclin D1 IHC, the Functional Genomics Center Zürich (FGCZ) for support with RNAseq and metabolomics, Michal Okoniewski (Scientific IT Services ETH) for help with bioinformatic analyses. Work in the Dechant, Peter, Curioni and Opitz laboratories is funded by the SNF, ETHZ and the Medical University of Zürich.

