## Supplementary Material, Figures and Methods for "Coincidence detection of mitogenic signals via cytosolic pH regulates Cyclin D1 expression"

contains

**Supplementary Figure legends**

**Materials and Methods**

**Supplementary References**

### Supplementary Figure legends:

#### Figure S1: NHE1 activity regulates pHc and promotes cell cycle progression through early G1

(**a-c**) FCS and glucose availability cooperatively regulate pHc. Cells were starved for 24h, stimulated with the indicated conditions and pHc was determined. Mean +/- S.E.M. of pHc (**a**), or the difference in pHc relative to the starved control (-/-) (**b-c**) was plotted as mean +/- S.E.M. alongside the histogram of single cell measurements. In the histograms, the median pHc was indicated. (**d**) NHE1 is the most abundant NHE localized to the plasma membrane. Relative expression levels of NHE1-9 were determined by qPCR (mean +/- S.E.M., N=3). Subcellular localization of the different NHE proteins is indicated. (**e-g**) NHE1 activity is required for RB phosphorylation. (**e**) The fraction of cells with RB phosphorylation at Ser780 from the experiment shown in Figure 1f is shown (mean +/- S.E.M., N= 3). (**f**) Cells treated as in Figure 1f were stained for $RB^{P\text{-}Ser807/811}$ and the fraction of cells with RB phosphorylation (mean +/- S.E.M., N= 3) (f) and representative images are shown. (**g**) Cells were subjected to siRNA against NHE1 and RB phosphorylation was determined after 72h by immunofluorescence or western blot. The fraction of RB positive cells (mean +/- S.E.M., N= 3) and representative images are shown. (**h-i**) RPE-1 cells were grown in media of different $NaHCO_3$ concentration to adjust extracellular pH to the indicated values and relative growth by MTT (mean +/S.E.M, N=2) (**h**), the fraction of cells with RB phosphorylation (mean +/S.E.M, N=2) and representative images (**i**) are shown.

**Figure S2: NHE activity regulates RB phosphorylation in different cell lines**

Cells treated as in Figure 1f were stained for RBP-Ser780 and the fraction of cells with RB phosphorylation was determined in the absence or presence of DMA for the indicated cell lines (mean +/- S.E.M., N= 3).

**Figure S3: pHc does not globally affect cellular metabolism.**

(**a-c**) RPE-1 cells were starved and the relative abundance of metabolites was determined 4h after stimulation with the indicated conditions as described in Materials and Methods. Selected metabolites from energy (**a**) and central carbon metabolism (**b**) are shown as mean +/- S.E.M.; N=4. P-values for all pairwise comparisons between conditions are listed in the Supplementary Table S7. (**c**) Unsupervised clustering and (**d**) PCA analysis were calculated using MATLAB (The MathWorks) from metabolite datasets generated using Progenesis QI software (Nonlinear Dynamics) to demonstrate the highly similar metabolic response upon stimulation in the presence or absence of DMA.

**Figure S4: Elevated pHc is required for early G1 progression.**

(**a-b**) Elevated pHc is required within the first 8h of the cell cycle. Extracts of cells as in Figure 1g were blotted for RB phosphorylation and Cyclin A expression as markers for G1 cell cycle progression. (**b**) Elevated pHc is required to drive cell cycle progression before initiation of S-phase. Cells were starved (-/-) and treated with FCS glucose for the indicated time points and BrdU incorporation was measured to follow DNA synthesis. Results of a representative experiment from three independent experiments with percentage of cells in G1, S and G2/M phase are shown. (**c-d**) pHc regulates a subset of FCS and glucose dependent genes required for G1 progression. Cells were starved for 24h, treated with FCS glucose in the presence or absence of DMA for 5h and analyzed by RNAseq. (**d**) Genes repressed upon starvation and DMA treatment and co-regulated genes in both conditions were analyzed by GO-term enrichment using ShinyGO (http://bioinformatics.sdstate.edu/go/[1]). Significantly enriched GO-terms were identified for the individual groups and plotted as -log10(FDR). (**e**) Venn diagram indicating the overlap of starvation and DMA-regulated genes. P-value demonstrating statistical significance of co-regulated genes based on a hypergeometric distribution is shown.

**Figure S5: Regulation of Cyclin D1 transcription requires CREB1/ATF1 and ETS1 transcription factors.**

(**a**-**c**) An elevated pHc is required for Cyclin D1 expression. (**a**) HFF-1 cells were starved and Cyclin D1 expression was determined by western blotting 6h after stimulation with the indicated conditions. (**b**) RPE-1 cells were grown and Cyclin D1 and phosphorylated RB was determined by western blotting upon treatment with DMA at the indicated time points. (**c**) RPE-1 cells were subjected to siRNA against NHE1 and Cyclin D1 abundance was determined after 72h by western blot (**d** and **e**) $CDK4^{Cyclin\ D1}$ activity is required for G1/S progression. Cells were treated as in Figure 1f and cell cycle progression was determined by (**d**) western blotting and (**e**) FACS analysis and upon treatment with the $CDK4^{Cyclin\ D1}$ inhibitor Ribociclib at the indicated times. (**f**) pHc regulates Cyclin D1 transcription rather than mRNA stability. Cells were treated with the indicated inhibitors for 4h and relative mRNA levels of Cyclin D1 were determined (mean +/- S.E.M., N=4) (**g**) CREB1, ATF1 and ETS1 are required for Cyclin D1 expression. Cells were transduced with Lentiviral particles expressing shRNA constructs against the indicated genes and expression of transcription factors and Cyclin D1 protein levels were determined by western blotting.

**Figure S6: Regulation of Cyclin D1 transcription requires p300/CBP**

(**a-b**) Reduced activity of p300/CBP decreases Cyclin D1 expression and acetylation of histone H3 at K27. (**a**) Cells were subjected to shRNA against the indicated transcription factors or co-activators and Histone acetylation at H3-K27 was determined by western blotting. (**b**) Cells were grown as in Figure 3b and Cyclin D1 mRNA levels were determined by 4h after stimulation with the indicated conditions by qPCR. (**c**) Cells were grown as in Figure 3a and the efficiency of shRNA mediated knock-down of p300 and CBP and Cyclin D1 protein levels were determined by western blotting. (**d**) DMA treatment and p300/CBP inhibition does not generally affect histone acetylation. Cells were grown as in Figure 3c and acetylation of Histone H3 at K9 was determined by western blotting.

**Figure S7: pHc and p300/CBP regulate a common transcriptional program.**

(**a** - **d**) Starvation, inhibition of NHE activity and p300/CBP activity trigger a similar gene expression program. Cells were grown as in Figure 1h and gene expression was determined by RNAseq. Volcano plots of differentially expressed genes upon stimulation with FCS glucose relative to (**a**) starvation, (**b**) DMA treatment and (**c**) C646 treatment is shown. Co-regulated genes under all conditions are indicated in red. (**d**) Venn diagram showing number of significantly higher expressed genes under the indicated conditions. (**e**) Unsupervised clustering of the normalized expression levels of co-regulated genes under all conditions.

**Figure S8: pHc is regulated by Akt activity and active glucose metabolism.**

(**a**) Cells as in Figure 3g were grown and relative biomass accumulation was scored by MTT 72h after DMA treatment. (mean +/- S.E.M.; N=6). (**b**) Cells as in Figure 3g were grown and RB phosphorylation was scored 24h after DMA treatment. (mean +/- S.E.M.; N=3). (**c**) CREB1 peptides interact with the KIX domain of CBP. A representative result of a pull-down of the KIX domain of CBP with biotinylated peptides is shown. (**d**) RPE-1 cells were starved, stimulated with the indicated conditions and changes in cytosolic pH (mean +/- S.E.M. of pooled single cell data, N=3) was determined. (**e**) RPE-1 cells were starved, stimulated with FCS and glucose in the presence of DMA and C646 and Akt-dependent phosphorylation of TSC2 was determined by western blotting. (**f**) Elevated pHc requires active glucose metabolism. pHc as determined in Figure 3j is plotted as mean +/- S.E.M. of pooled single cell data of at least 3 independent experiments.

**Figure S9: NHE1 activity is critical for cell cycle progression in PDAC cell lines.** (**a**) Cells of the indicated genotype expressing pHluorin were starved for 24h and pHc was determined 30 min following stimulation with the indicated conditions. (**b**) Cells as in (**a**) were starved, treated with DMA for 6h and Cyclin D1 and RB phosphorylation were determined by western blotting. (**c**) Cells were subjected to siRNA against NHE1 and Cyclin D1, RB phosphorylation and NHE1 abundance were determined by western blotting. (**d**) Cyclin D1, but not NHE1 expression correlates with survival in PDAC cell lines. Data obtained from TCGA were independently analyzed for a correlation of Cyclin D1 and NHE1 mRNA and survival and Kaplan Meier analysis was performed.

**Figure S10: MPMs may be caused by hyperactivation of the NHE1/Cyclin D1 axis** (**a**) MPMs are associated with high $CDK4^{Cyclin\ D}$ activity. Percentage of patients with the indicated mutations was determined from 2 patient cohorts as described in Materials and Methods. Only mutations in genes linked to the Cyclin D/Rb pathway are shown. (**b**) Cells were grown and treated as in Figure 4c and Cyclin D1 protein abundance was analyzed by western blot. (**c**) Cells were grown and phosphorylation of RB was analyzed upon DMA treatment (mean +/- SEM, N=3). (**d**) Cells subjected to siRNA mediated knockdown of NHE1 and Cyclin D1 and NHE1 protein levels were analyzed by western blot. (**e**) NHE1 is the most abundant NHE isoform in MPM cell lines. Steady state mRNA levels of the indicated genes were determined by qPCR and plotted as mean +/- SEM relative to NHE1 (N=3).

**Figure S11: NHE1 expression correlates with Cyclin D1 expression and survival in MPM patients.**

(**a**) Representative images of histological sections stained as in Figure 4f. (**b**) Kaplan Meier analysis of survival data based on high or low NHE1 expression from the TCGA-Meso cohort. (**c**) Cyclin D1 expression for patient cohorts with high and low NHE1 expression as determined in (**b**).

### Materials and Methods:

#### Standard molecular biology methods:

Standard manipulations were used for generating plasmids and lentiviral particles and for culturing cells. Plasmids used in this study are listed in Table S1. Unless otherwise stated, RPE-1 cells were maintained in DMEM media supplemented with FCS (10% final concentration) and Penicillin Streptavidin-Glutamine (PSG; Gibco, 1% final concentration), MPM cell lines were maintained in RPMI containing FCS (10% final), Glucose (5 mg/ml) and PSG. For starvation, cells were washed once with RPMI medium (Gibco, w/o glucose, FCS and PSG) and incubated in the same medium for 24h. Cells were stimulated with RPMI medium containing FCS (10% final) and Glucose (5 mg/ml) as indicated. For adjusting extracellular pH RPMI powder (Sigma-Aldrich) was reconstituted according to manufacturer's instructions and $NaHCO_3$ was adjusted using a 7.5% $NaHCO_3$ solution (ThermoFisher) to a final concentration of 44mM or 5 mM, respectively. Western blotting was performed using standard methods. Antibodies used for detection of specific proteins are listed in table S2 and proteins were visualized using a Fusion Imaging System (Witec AG). Treatments with pharmacological inhibitors were performed as summarized in Table S3. FACS analysis and BrdU incorporation measurements were performed as described previously[2]. Details on oligo sequences for qPCR, RNAi and ChIP are listed in Supplementary Tables S4-S6.

#### pH measurements:

For measurement of cytosolic pH, 90000 RPE cells expressing SEpHluorin were seeded in an 8-well ibidi coverslip (ibidi) coated with poly-lysine-D (0.1 mg/ml in $H_20$, Sigma-Aldrich, 5 min at RT) in 200 $\mu$l of growth medium. Cells were grown for at least 24h before being starved for serum and glucose for 24h. Cell were stimulated with 250 $\mu$l fresh medium containing glucose and FCS and inhibitors as indicated and incubated for 1h at 37C with 5% CO2. For all cytosolic pH measurements with the SEpHluorin constructs, images were captured at an inverted Ti-Elcipse (Nikon) microscope upon excitation with CFP (436 +/- 20 nm) and YFP (500 +/- 20 nm) excitation filters. Emission was recorded with a YFP emission filter (550 +/- 30 nm) filter and the ratio of intensities in these channels was quantified for a small rectangular region representing the cytosol following background correction in at least 30 individual cells per condition and experiment using ImageJ (http://rsb.info.nih.gov/ij/).

For generating a pH standard curve, cells were washed briefly twice with a $K_2HPO_4$ - $KH_2PO_4$ buffer adjusted to a pH of either 7.0, 7.2, 7.4, or 7.6, before cells were incubated in 250 $\mu$l of the buffers containing 10 $\mu$M nigericin (Thermo Fisher Scientific) for 10 to 20 min at 37 C without $CO_2$ control before imaging. Images were quantified as before and calibration curves were fit to determine the pH values for individual cells using Excel.

For measurement of cytosolic pH in MPM cell lines, cells were grown in 8-well ibidi coverslips (ibidi), washed once with PBS and incubated with the pH-sensitive dye SNARF-4 (Invitrogen, 10$\mu$M in PBS containing 2mM glucose) for 30 minutes. Then cells were incubated with fresh growth medium in the presence of 5% CO2 for 1h and imaged using filter sets appropriate for RFP (excitation: 560 +/- 40 nm, emission: 630 +/- 75 nm) and GFP (470 +/- 40 nm, 525 +/- 50 nm). Image analysis and calibration were performed as described above.

**MTT assay:**

MTT (Pierce) assays were performed essentially as described by the manufacturer. In brief, 3000-5000 cells were seeded into 96-well plates and incubated for 3 days. MTT was added for 2h before reactions were stopped and dye formation was measured at OD550.

**qPCR analysis**

Total RNA was extracted from cells using the RNeasy kit (Quiagen). RNA was reverse transcribed using random hexanucleotides and SuperScript II polymerase (Roche) and relative abundance of specific mRNAs was determined on a Roche Lightcycler using the SYBRgreen method. GAPDH and β-Actin were used as reference genes for RPE-1 cells. For the determination of optimal reference genes for malignant pleural mesotheliomas, RNAseq data were downloaded from the Broad Institute (RPKM, release 20180502, http://www.broadinstitute.org/ccle, [3]). The coefficient of variation (CV) of expression levels of genes detected in all cell lines derived from malignant pleural mesotheliomas of the epitheloid subtype was determined. Top hits with lowest CV were tested for suitability for qPCR analysis and PPP4C and SPOPL were chosen as reference genes.

**Chromatin Immunoprecipitation (ChIP):**

Chromatin Immunoprecipitation was performed using the ChIP kit from CellSignaling technology according to manufactures instructions. In brief, cells from 2 15 cm dishes were cross linked using formaldehyde for 15 minutes and harvested. Cells were lysed and nuclei were isolated by centrifugation. DNA was fragmented by Nuclease digestion and nuclei were disrupted by brief sonication. Cleared extracts were subjected for immunoprecipitation using P-CREB1, p300 and IgG antibodies as control. After reversal of the cross-link, DNA was isolated and analyzed by qPCR using the Cyber green method on a Lightcycler (Roche). Primers used for amplification of genomic fragments corresponding to the CRE-element within the Cyclin D1 promoter (-50 nt) and a distal site within the ORF (+8000 nt downstream of the ATG) are listed in Table S6 (sequences from [4]q).

**Subcellular protein fractionation:**

Subcellular protein fractionation and isolation of chromatin fractions was performed using the Subcellular Protein Fractionation Kit for Cultured Cells kit from Thermo Fisher following the instructions of the manufacturer. Soluble nuclear and chromatin bound nuclear extracts were loaded on an SDS-page gel and subjected to western blotting.

**Determination of IC50 values:**

For the determination of IC50 values, 3000-5000 cells were seeded in 96-well plates and grown in the presence of 2-fold serial dilutions of the indicated drugs or drug vehicle alone and biomass was determined after 3 days using MTT. IC50 values were determined using Excel or Prism software.

**Metabolomics:**

**Sample preparation for LC-MS analysis**

For metabolite extraction, cells were incubated with a Methanol:Chloroform:H2O (60:20:20 %Vol) mixture with shaking for 3 hours at 4 °C, before the supernatant was cleared by centrifugation to remove debris and dried. The resulting pellets were stored at –80 °C until reconstitution in 20 $\mu$L water (MS grade) and dilution with 80 $\mu$L injection buffer. The dilution was vortexed and centrifuged (16,000 x g, 4 °C, 15 min). 50 µL of the supernatant was transferred to a glass vial with narrowed bottom (Total Recovery Vials, Waters) for LC-MS injection. In addition, method blanks, QC standards, and pooled samples were prepared in the same way to serve as quality control for the measurements. Injection buffer was composed of Acetonitrile:Methanol:5mM ammonium acetate (90:9:1 %Vol).

**LC-MS analysis**

Metabolites were separated on a nanoAcquity UPLC (Waters) equipped with a BEH Amide capillary column (150 μm x50mm, 1.7 μm particle size, Waters), applying a gradient of 5mM ammonium acetate in water (A) and 5mM ammonium acetate in acetonitrile (B) from 5% A to 50% A over 12 min. The injection volume was 1 μL. The flow rate was adjusted over the gradient from 3 to 2 μl/min.

The UPLC was coupled to a Synapt G2Si mass spectrometer (Waters) by a nanoESI source. MS1 (molecular ion) and MS2 (fragment) data were acquired using negative polarization and MSE over a mass range of 50 to 1200 m/z at MS1 and MS2 resolution of >20'000.

**Targeted Data analysis**

Selected metabolites within the energy and central carbon metabolism were quantified by the area under the peak (AUP) of the MS1 extracted ion chromatogram (EIC) of the respective [M-H]- ion extracted from raw data by using MassLynx v4.2 software (Waters). For all targets reference samples were used to locate and verify the correct peaks on the EIC of the samples by retention time and MS2 fragment information. AUP values from individual samples were normalized by the area of the total ion count (TIC) of each sample to account for variation of biomass in the samples between experiments and normalized relative to the control condition. Note that G6P/ F6P, and Citrate/Isocitrate were quantified as the sum of both metabolites due to limited peak separation.

**Untargeted Metabolomics Data analysis**

For analysis of global metabolic changes, data sets were evaluated in an untargeted fashion with Progenesis QI software (Nonlinear Dynamics), which aligns the ion intensity maps based on a reference data set, followed by peak picking on an aggregated ion intensity map. Heat maps and PCA analysis were calculated w/o identification of the detected peaks following normalization of intensities across samples using MATLAB.

**RNAseq and data analysis:**

**RNA isolation and library prep:**

For RNAseq experiments, total RNA was isolated using the RNeasy kit (Quiagen). RNA concentration was determined by a Qubit® (1.0) Fluorometer (Life Technologies, California, USA) and quality of RNA was assessed using a Fragment Analyzer (Agilent, Santa Clara, California, USA). Only samples with a 260 nm/280 nm ratio between 1.8–2.1 and a 28S/18S ratio within 1.5–2 were further processed. RNA was diluted 2-fold and 10 $\mu$l of RNA was supplemented with 3$\mu$l of ERCC ExFold RNA Spike-In Mix 1 (dilution 1:100) before RNA concentration was adjusted to 50ng/$\mu$l and submitted for library preparation using the TruSeq Stranded mRNA kit (Illumina, Inc, California, USA). In brief, total RNA samples (100-1000 ng) were polyA enriched and then reverse-transcribed into double-stranded cDNA. The cDNA samples were fragmented, end-repaired and adenylated before ligation of TruSeq adapters containing unique dual indices (UDI) for multiplexing. Fragments containing TruSeq adapters on both ends were selectively enriched with PCR. The quality and quantity of the enriched libraries were validated using a Qubit® (1.0) Fluorometer and the Fragment Analyzer (Agilent, Santa Clara, California, USA). The product is a smear with an average fragment size of approximately 260 bp. The libraries were normalized to 10nM in Tris-Cl 10 mM, pH8.5 with 0.1% Tween 20.

**Cluster Generation and Sequencing**

Cluster generation and sequencing was performed on a Novaseq 6000 sequencer (Illumina, Inc, California, USA) according to manufacturer's instructions. Sequencing was done at 2x 150 bp (paired end).

**Data analysis**

Raw reads were first cleaned by removing adapter sequences, trimming low quality ends, and filtering reads with low quality (phred quality <20) using Trimmomatic (Version 0.36)[5]. Read alignment was done using STAR (v2.7.0e)[6]. As reference we used the Ensembl genome build GRCh38.p10 with the gene annotations downloaded on 2017-05-31 from Ensembl (release 89) and extended by the ERCC spike-in sequences. STAR alignment options were as follows:

--outFilterType BySJout; --outFilterMatchNmin 30; --outFilterMismatchNmax 10;
--outFilterMismatchNoverLmax 0.05; --alignSJDBoverhangMin 1;
--alignSJoverhangMin 8; --alignIntronMax 100000; --alignMatesGapMax 100000;
--outFilterMultimapNmax 50

Sequence pseudo alignment of high-quality reads to the Human reference genome (build GRCh38.p10) and quantification of gene level expression was carried out using Kallisto (Version 0.44)[7]. Raw reads per gene were extracted for the ERCC spike-in sequences for each individual sample and fitted using a linear regression model, which was applied to raw reads of genes. To detect differentially expressed genes we applied the Bioconductor package EdgeR[8] (R version: 3.5.1, EdgeR version: 3.24.3. Differential expression was assessed using a genewise negative binomial generalized linear model with biological questions coded as contrasts. Genes showing altered expression with adjusted (Benjamini and Hochberg method) p-value at FDR < 0.05 and an absolute log2(fold-change) of >0.5 were considered differentially expressed.

**Enrichment analysis:**

For testing the enrichment of E2F targets in the dataset in Figure 1, experimentally verified E2F1 targets in RPE1-hTERT cells were obtained from the NCBI Sequence read archive (https://www.ncbi.nlm.nih.gov/sra/?term=SRX1036433) via ChipAtlas.org.[9] A two-tailed unpaired t-test was applied on the log2(fold-changes) for E2F targets vs. all detected genes to assess statistical significance and confirmed by calculating the probability of enrichment of differentially expressed E2F targets (repressed by DMA, log2(fold-change) > 0.5) relative to all verified E2F1 targets in the dataset of Figure 1 applying a hypergeometric distribution model ($P<2.5x10^{-8}$).

Enrichment of process networks as well as transcription factor networks for differentially expressed genes was determined using MetaCore from Clarivate Analytics (MetaCore+MetaDrug® version 19.3 build 69800, http://portal.genego.com).

**Bioinformatics:**

**Expression analysis:**

For expression analysis of NHE1 and Cyclin D1 in different cancer cell-lines, RNAseq data were downloaded from the Broad Institute (RPKM, release 20180502, http://www.broadinstitute.org/ccle, [3]) and Pearson correlation coefficients were determined between SLC9A1 (NHE1) and CCND1 (Cyclin D1) expression for data sets of individual cancer types using MATLAB.

**Survival curves**

For calculation of survival curves, data from the Mesothelioma cohort (TCGA-MESO) or det PDAC cohort (TCGA-PAAD) were downloaded from the TCGA database (https://www.cancer.gov/tcga) and analyzed using custom made, MATLAB based scripts. P-values, median survival times and hazard ratios were determined based on a Mantel-Cox log rank test using the MATLAB based script MatSurf.m.[10]

**Analysis of genetic alterations in MPMs**

For mutational analysis of potential driver mutations in Mesothelioma patients, genomic data were obtained from the cBioPortal (http://www.cbioportal.org, [11,12]), and data of three studies of Mesotheliomas were pooled (TGCA pan cancer atlas, https://portal.gdc.cancer.gov, and [13]).

**Luciferase assays**

RPE-1 cells were seeded at 3 x $10^5$ per well of a 6-well plate and allowed to adhere overnight. $24^h$ post-transfection cells were washed with 1x PBS and starved for FCS and glucose for $24^h$. Luciferase assays were performed 6h after stimulation with the indicated conditions using the Pierce™ Firefly Luciferase Glow Assay Kit according to the manufacturer's instructions.

For Cyclin D1 promoter mutant constructs, cells were transfected with either fragments of the wild-type Cyclin D1 promoter fused to Luciferase or promoter mutants lacking transcription factor binding sites (ΔCREB, ΔEtsA, ΔEtsB, ΔEtsC, ΔAP1) using the Lipofectamine™ 3000 Transfection Reagent. Luciferase assays were performed 48h post transfection using the Pierce™ Firefly Luciferase Glow Assay Kit according to the manufacturer's instructions.

For GAL4-CREB1 based assays, cells were co-transfected with either 5xGAL4-TATA-luciferase (2500ng) and GAL4-CREB1 (2500ng) or 5xGAL4-TATA-luciferase and GAL4-CREB1$^{DIEDML}$ using the Lipofectamine™ 3000 Transfection Reagent. 12 hours post-transfection 20 000 cells per well were plated onto a 96-well plate (24 wells per condition). $24^h$ post-transfection cells were washed with 1x PBS and starved for FCS and glucose for $24^h$ and Luciferase assays were performed as above.

##### Protein expression and binding assays

The KIX domain of CBP was expressed from a pET21-a based vector[14] in *E.coli* (BL21) following IPTG induction (0.4 mM) at 15°C overnight. Cells were harvested and washed once in PBS before, resuspended in KIX-100 buffer (20 mM HEPES, 100 mM KCl, 0.2 mM EDTA, 5 mM MgCl2, 0.1% NP40, 20% Glycerol, 100$\mu$g/ml BSA, Roche protease inhibitors and PMSF 1$\mu$M, pH 6.9) and lysed using cryo-milling. Extracts were cleared by centrifugation (20min at 48000g) and stored at -80°C until further use. For peptide binding assays, 20$\mu$l of Strep-beads (Strep-Tactin® Sepharose®, IBA Lifesciences) were incubated with 10$\mu$g of biotinylated peptides dissolved in 200$\mu$l KIX-buffer for 30 min at 4°C and washed 3x with KIX buffer. Then, beads were incubated with 200 $\mu$l bacterial extract containing KIX domain for 45min at 4°C, washed 3x with KIX buffer, and washed once with KIX-100 buffer adjusted to different pH using NaOH or HCl, respectively. Beads were incubated with adjusted KIX-100 buffers for 15 min at RT, and washed 3x. Proteins and peptides were eluted using 50$\mu$l preheated sample buffer containing 8M urea and analyzed by SDS-page using 4-12% NuPAGE gels (Invitrogen) in MES buffer according to manufacturer's instructions.

##### Histology

**Patient cohort:**

To score expression of NHE1 and Cyclin D1, respectively, we retrospectively evaluated NHE1 and Cyclin D1 expression by immunohistochemical (IHC) analysis in resected specimens (in form of a tissue microarray) in patients with malignant pleural mesothelioma before receiving or not chemotherapy after surgical resection. 96 patients (9 female (9%) and 87 males (91%)) were included and 2 specimen per patient were analyzed. The histological subtype was epithelioid in 70 patients (73%), biphastic in 16 patients (17%), sarcomatoid in 5 patients (5%) and undefined in 5 patients (5%).

12 patients (13%) received Cisplatin, 29 (30 %) a combination of Cisplatin and Gemcitabine, 2 (2%) a combination of Carboplatin and Permetrexed, 31 (32%) a combination of Pemetrexed and Cisplatin and 22 (23%) no therapy.

All patients have given informed consent and the study was approved by the Ethical Committee Zürich (EK-ZH 2012-0094 and EK-ZH 2018-01919).

**Staining and analysis:**

For staining, paraffin embedded tumors were cut into 2 $\mu$m cuts, mounted on a glass slide and dried overnight. The next day tumor sections were deparaffinised and rehydrated, antigen retrieval was performed for 40min using the Epitope Retrieval Solution 2 (Leica). Slides were stained with a monoclonal anti-Cyclin D1 (Labvision RM-9104-S, clone SP4, 1:40) or anti-SLC9A1 (Sigma Aldrich (HPA052891) 1:100). The immunohistochemical staining was performed using Leica Bond III automated system including Polymer Refine DAB Kit.

A score of 0 (low) or 1 (high) was assigned to individual tumor sections and averaged over all spots from the same patient to yield a mean expression score. Tumors were grouped based on this analysis for NHE1 in tumors in low NHE1 expressing (score equal to, or lower than 0.5) and high NHE1 expressing tumors (score greater than 0.5). Based on these groups, the score of Cyclin D1 was plotted as the mean +/-S.E.M., and Kaplan Meier analysis was performed using Prism applying the log-rank test.

#### Statistical analysis

Unless otherwise stated, the data are represented as mean +/- S.E.M. of at least three independent replicates and two-tailed Student's t-tests were calculated to assess statistical significance.

For pH measurements, single cell data of at least three independent experiments were pooled represented as mean +/- S.E.M. Histograms of single cell data were calculated using MATLAB to indicate the difference in pHc relative to the control condition. Mean +/- S.E.M. are displayed next to the histograms and the median pH value is indicated together with the histograms.

For quantification of mRNA levels using RNAseq, relative expression levels of the gene of interest were quantified relative to two different reference genes and the obtained fold changes for each reference genes were averaged for each experiment and normalized to the control condition. Data are represented as mean +/- S.E.M. of at least three independent replicates.

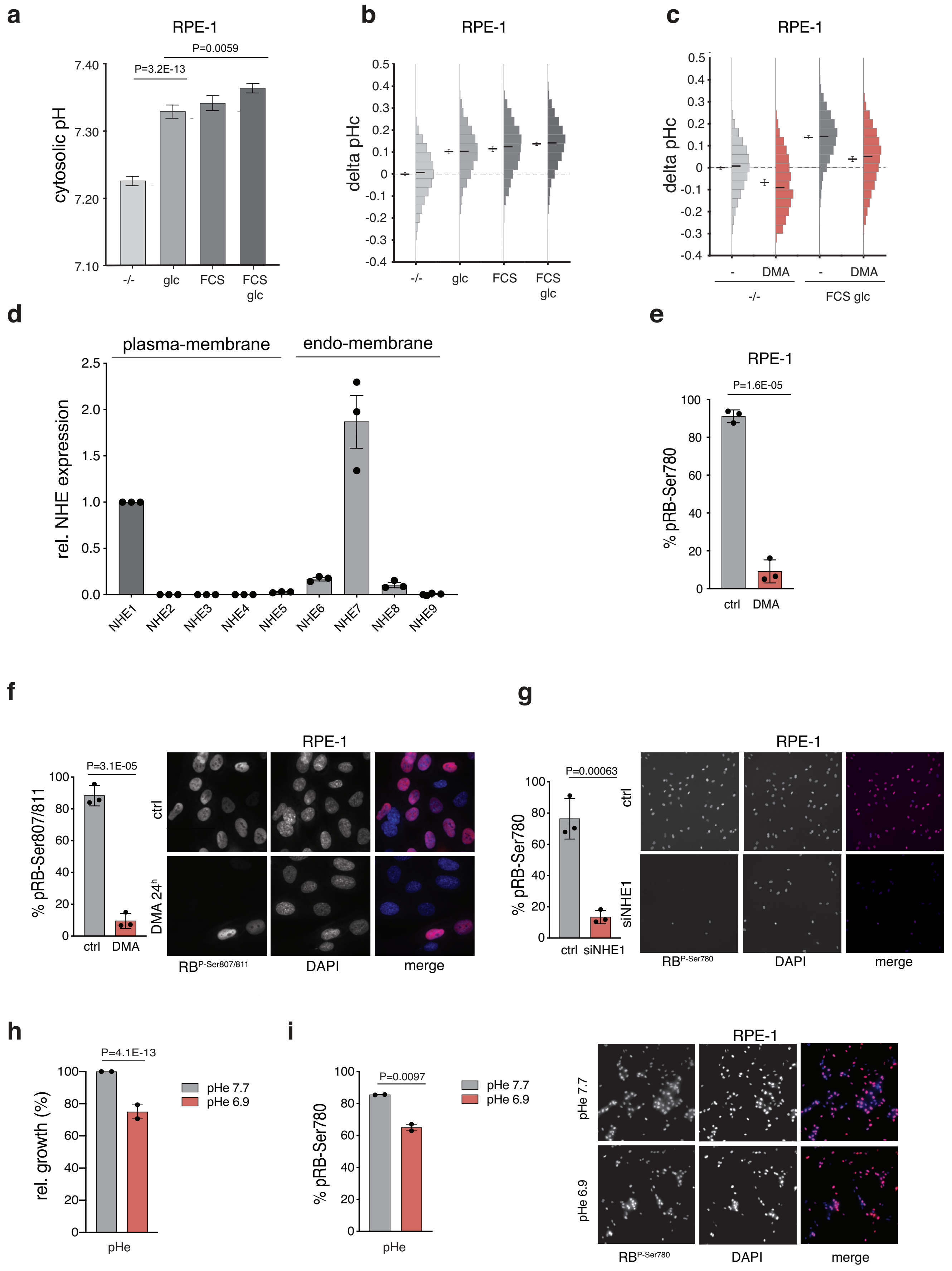


Figure S1a: NHE1 activity regulates cytosolic pH and promotes cell cycle progression through early G1

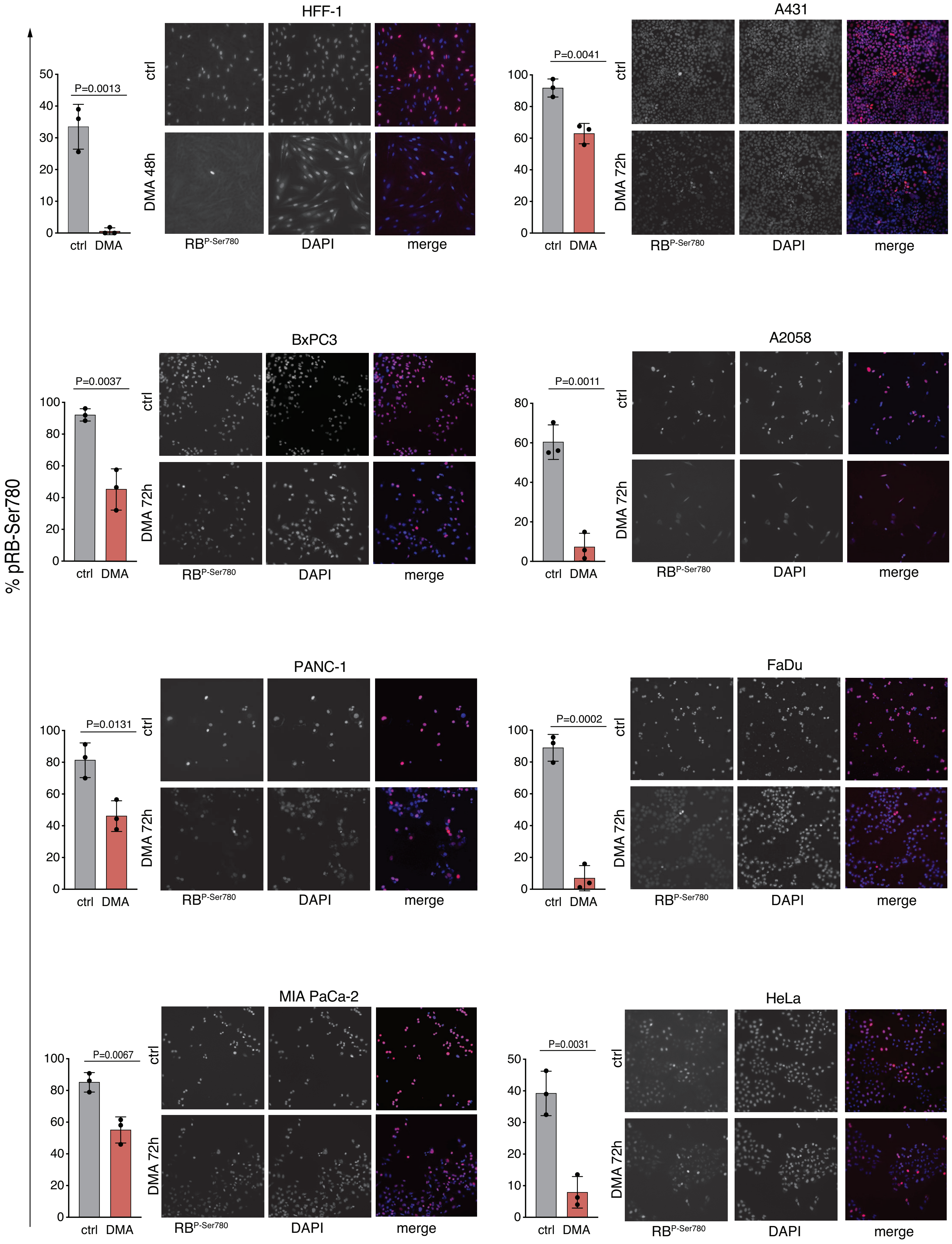


Figure S2: NHE1 activity regulates RB phosphorylation in different cell lines

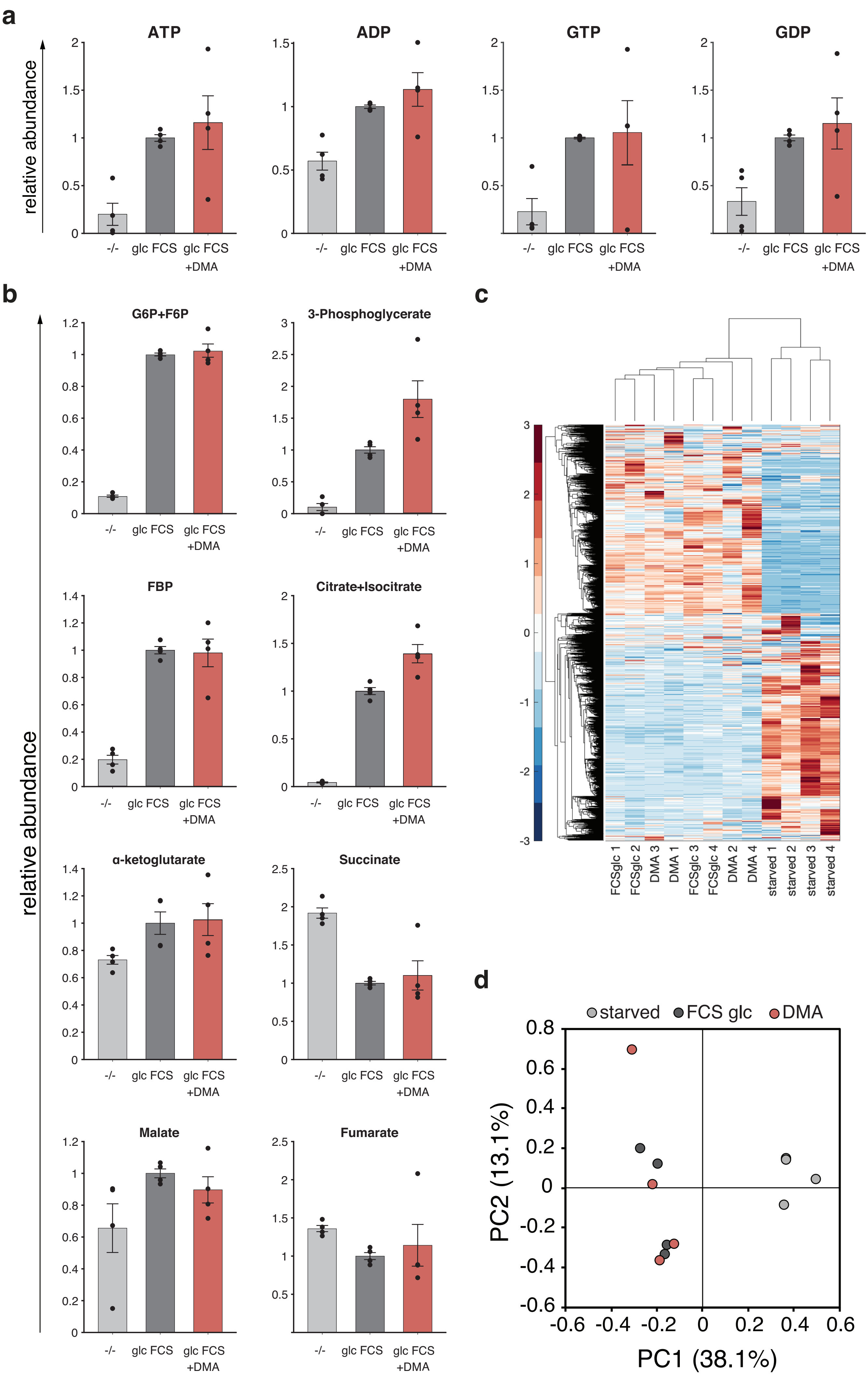


Figure S3: pHc does not globally affect cellular metabolism

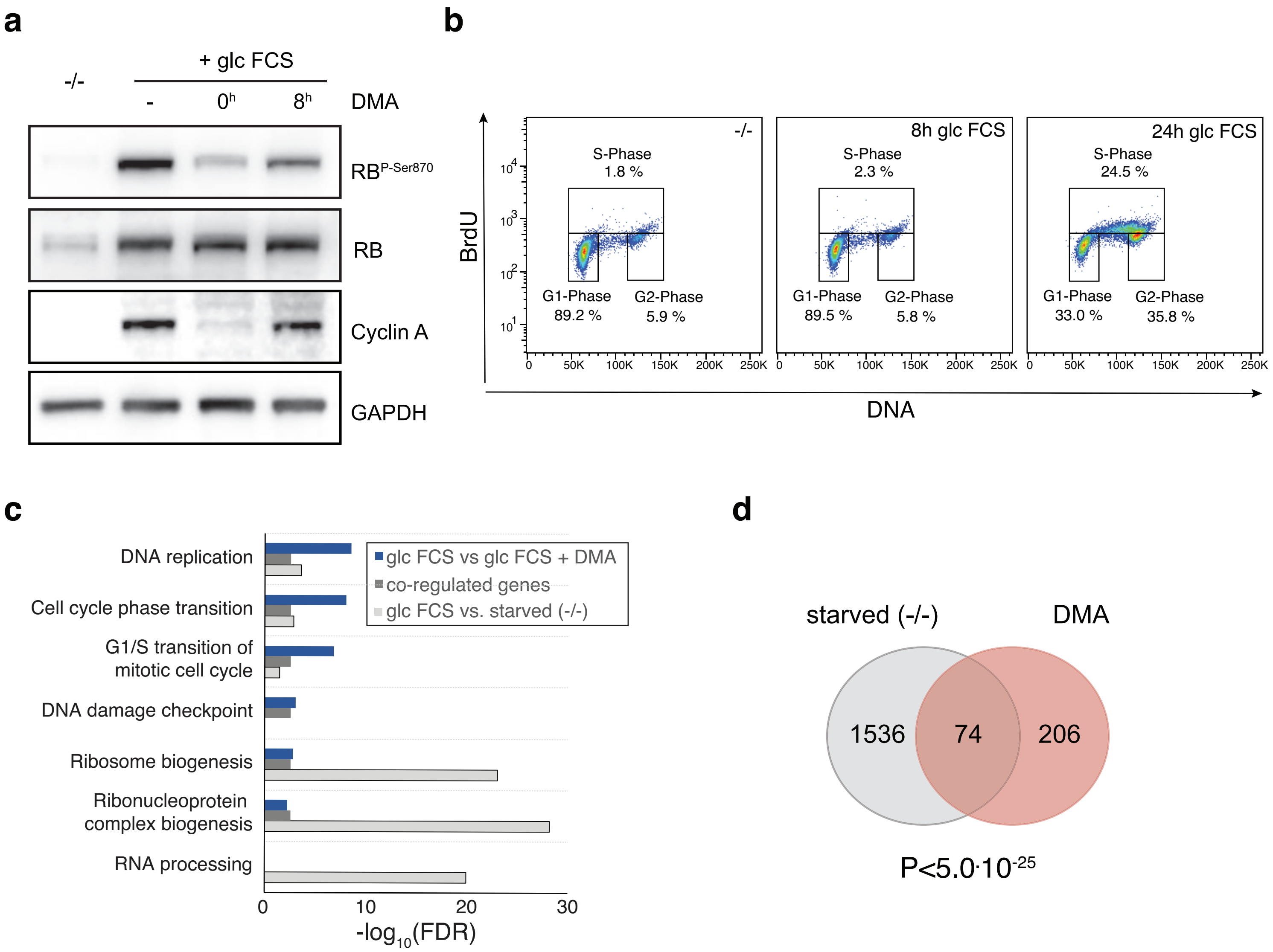


Figure S4: Elevated pHc is required for early G1 progression

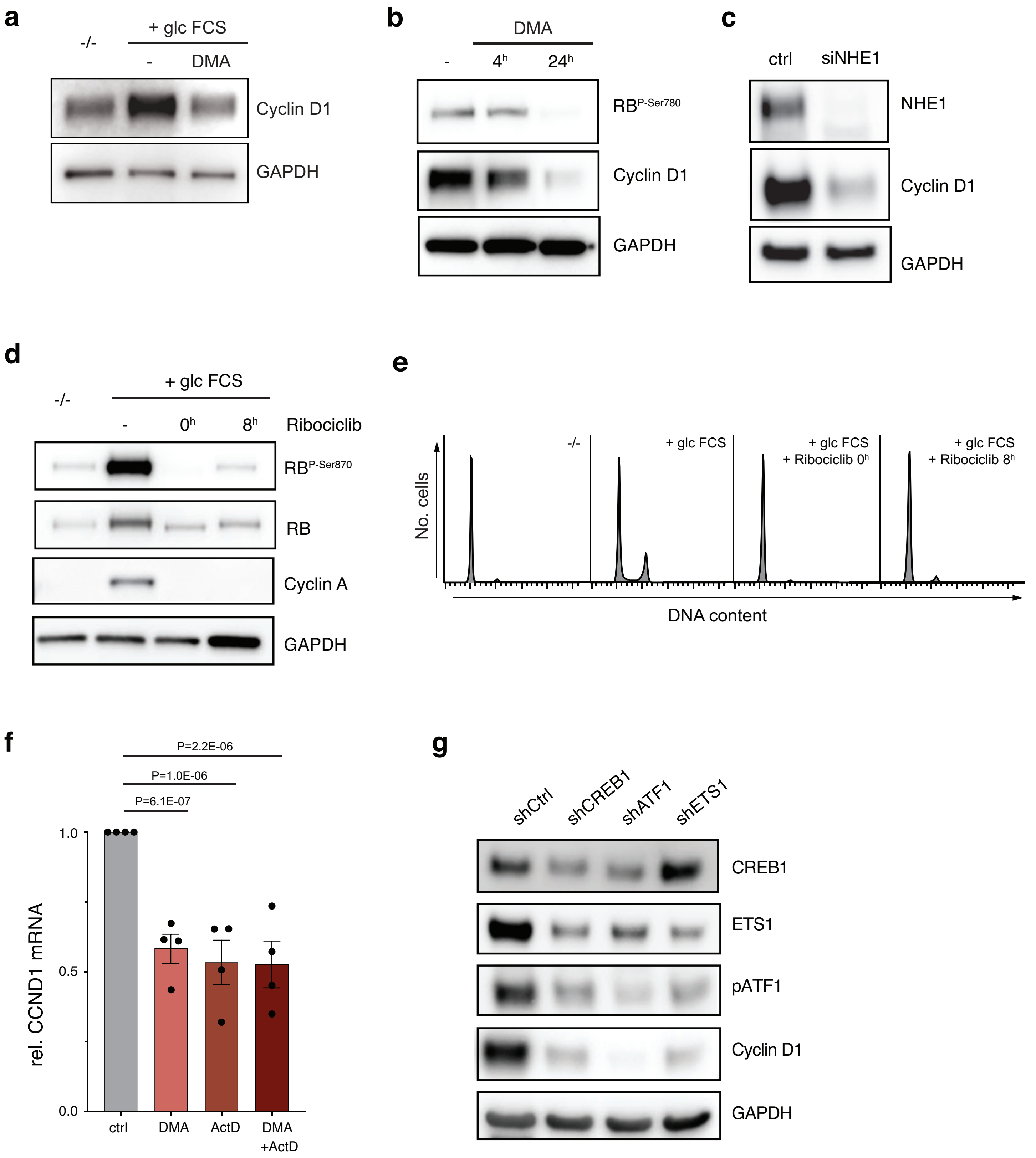


Figure S5: Regulation of Cyclin D1 transcription requires CREB1/ATF1 and ETS1 transcription factors

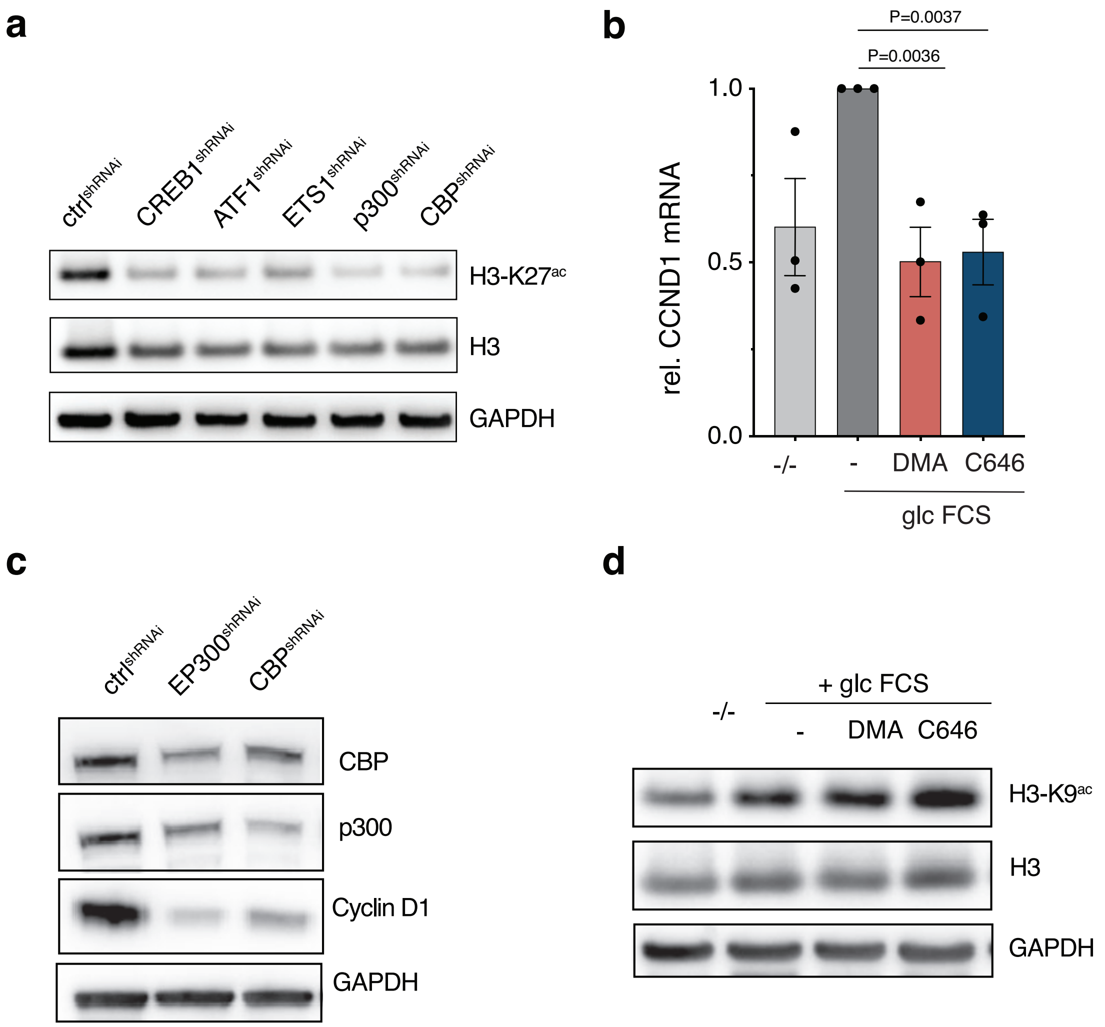


Figure S6: Regulation of Cyclin D1 transcription requires p300/CBP

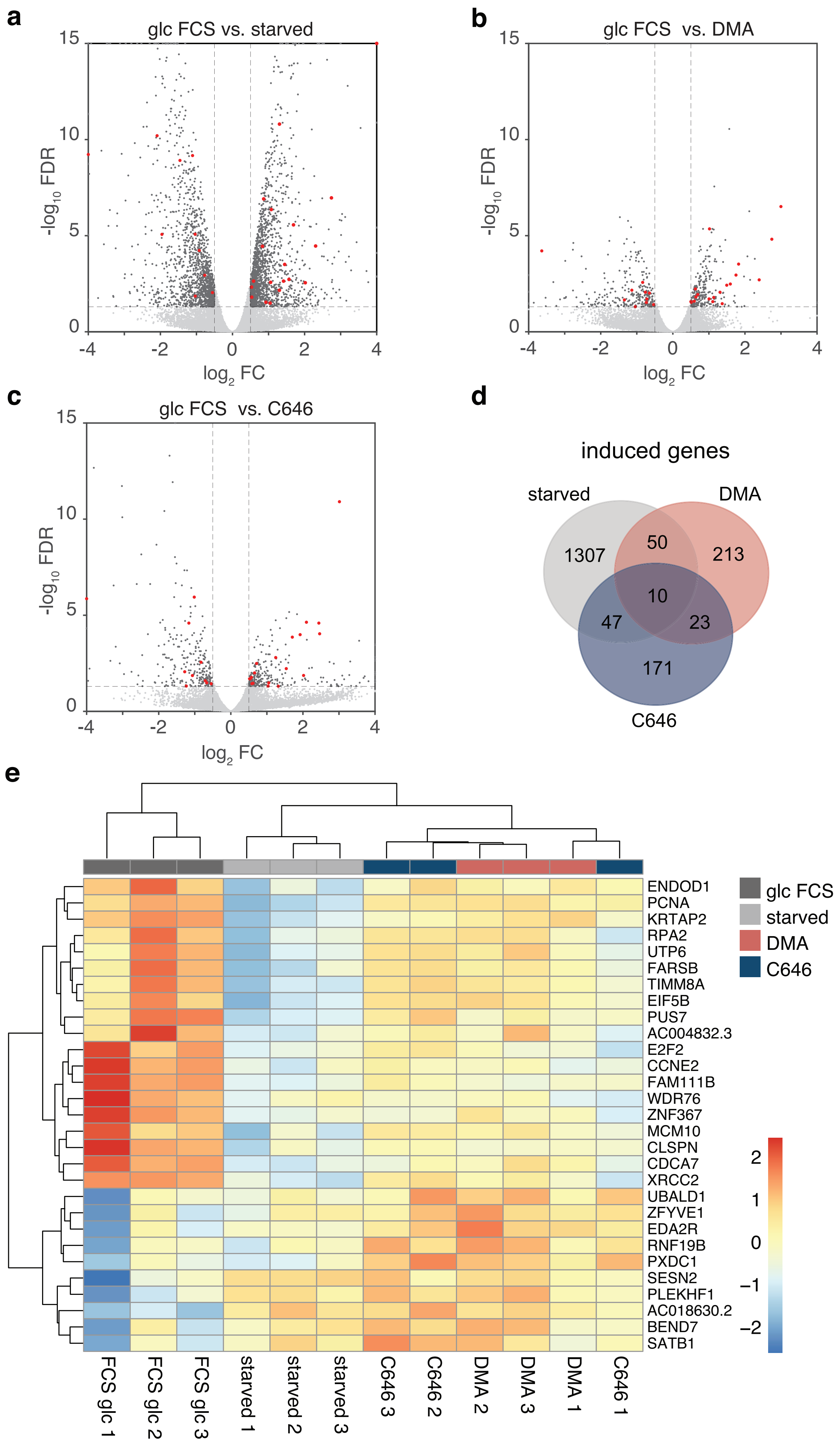


Figure S7: pHc and p300/CBP regulate a common transcriptional program

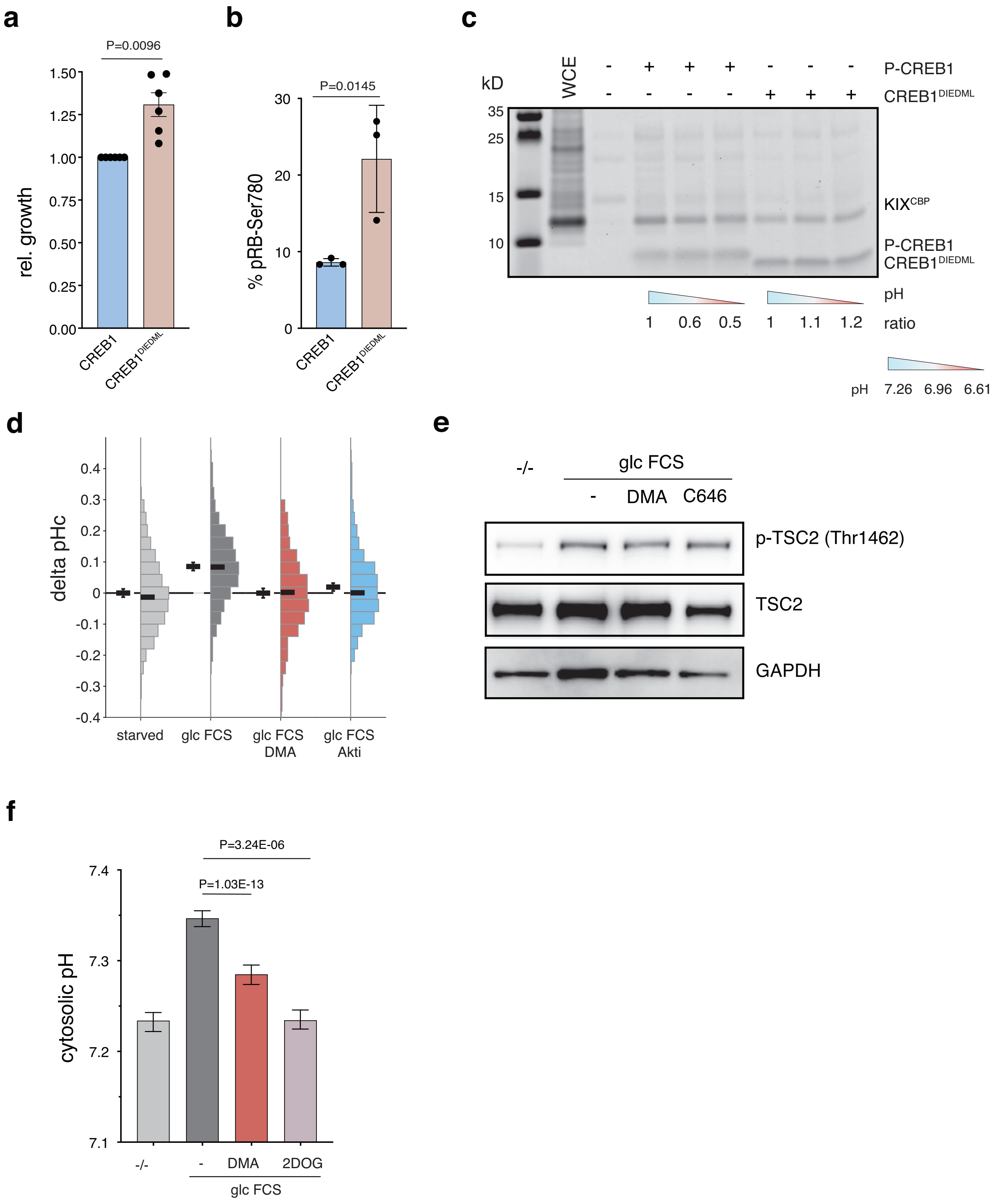


Figure S8: pHc is regulated by Akt activity and active glucose metabolism.

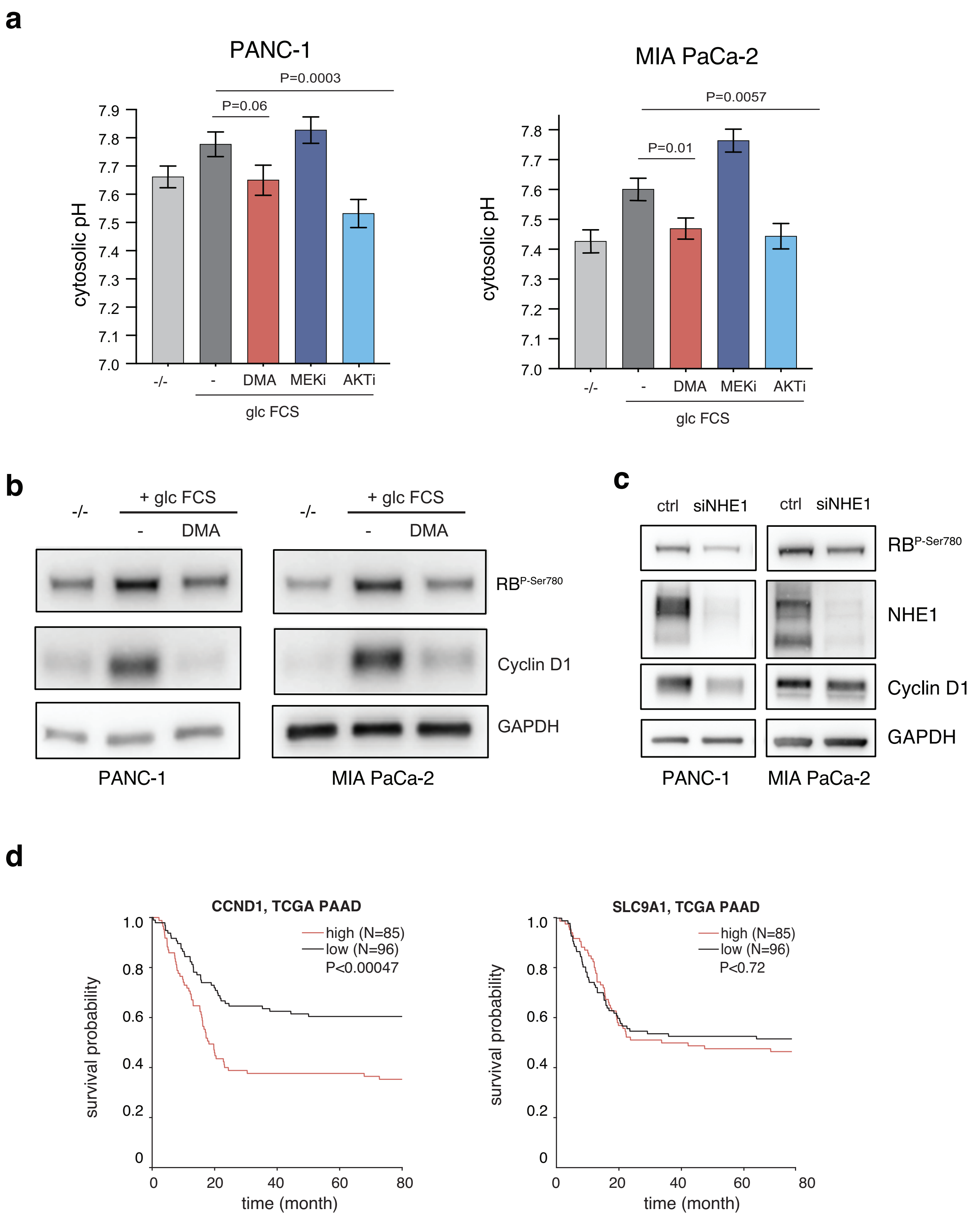


Figure S9: NHE1 activity is critical for cell cycle progression in PDAC cell lines

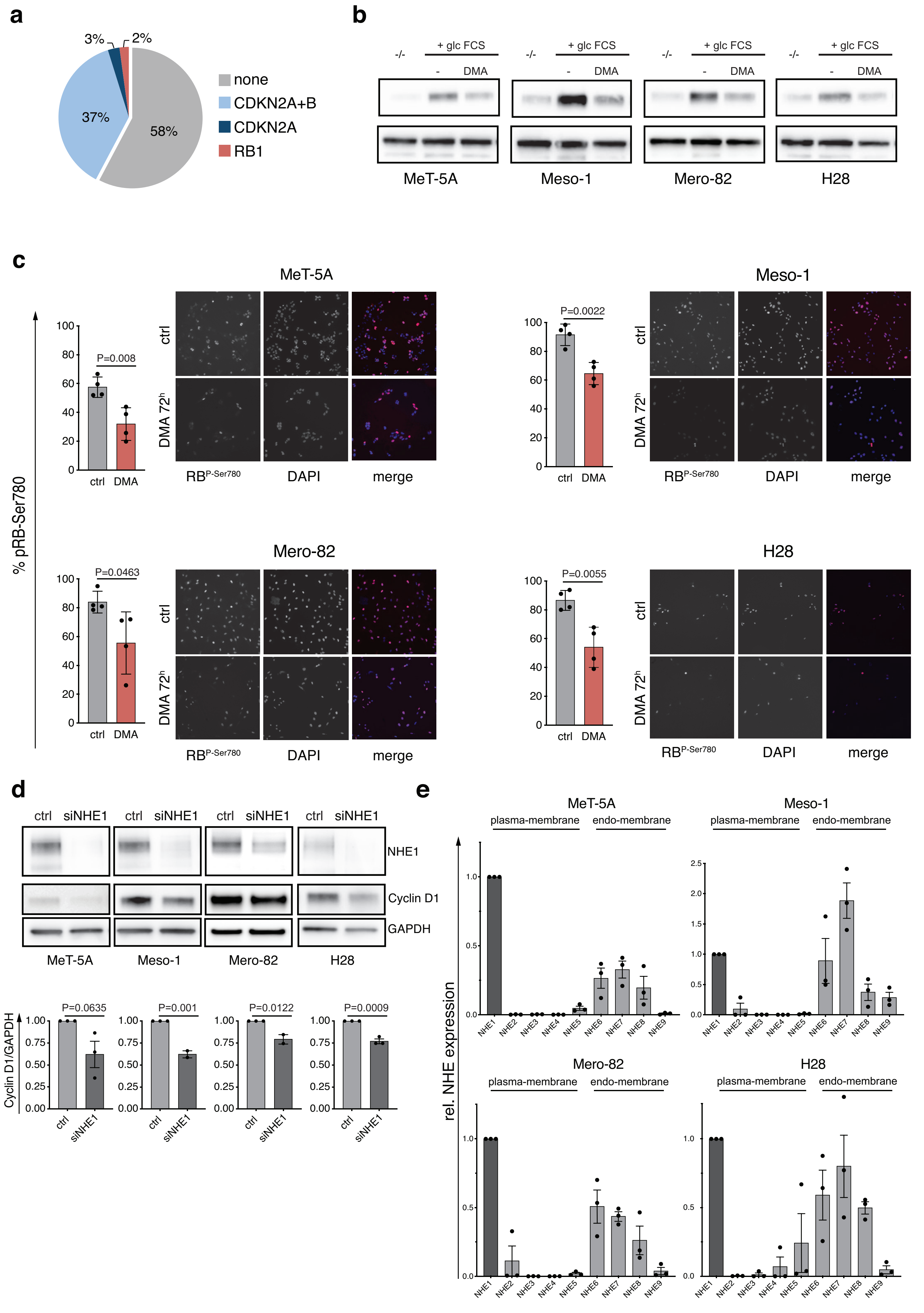


Figure S10: MPMs may be caused by hyperactivation of the NHE1/Cyclin D1 axis

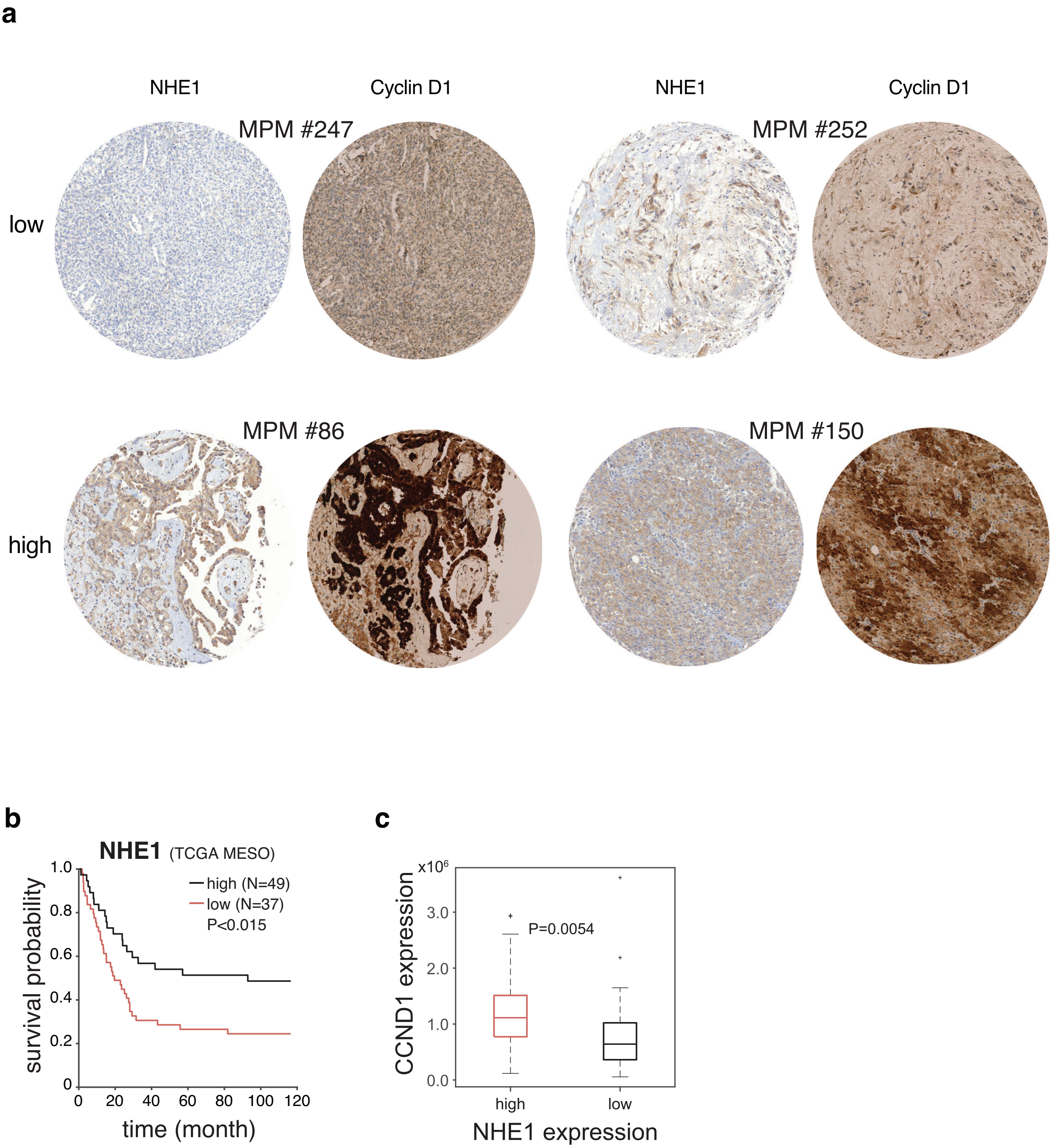


Figure S11: NHE1 expression correlates with Cyclin D1 expression and survival in MPM patients
